# AliSim: A Fast and Versatile Phylogenetic Sequence Simulator For the Genomic Era

**DOI:** 10.1101/2021.12.16.472905

**Authors:** Nhan Ly-Trong, Suha Naser-Khdour, Robert Lanfear, Bui Quang Minh

## Abstract

Sequence simulators play an important role in phylogenetics. Simulated data has many applications, such as evaluating the performance of different methods, hypothesis testing with parametric bootstraps, and, more recently, generating data for training machine-learning applications. Many sequence simulation programs exist, but the most feature-rich programs tend to be rather slow, and the fastest programs tend to be feature-poor. Here, we introduce AliSim, a new tool that can efficiently simulate biologically realistic alignments under a large range of complex evolutionary models. To achieve high performance across a wide range of simulation conditions, AliSim implements an adaptive approach that combines the commonly-used rate matrix and probability matrix approach. AliSim takes 1.3 hours and 1.3 GB RAM to simulate alignments with one million sequences or sites, while popular software Seq-Gen, Dawg, and INDELible require two to five hours and 50 to 500 GB of RAM. We provide AliSim as an extension of the IQ-TREE software version 2.2, freely available at www.iqtree.org, and a comprehensive user tutorial at http://www.iqtree.org/doc/AliSim.

## Introduction

Simulating a multiple sequence alignment (MSA) plays a vital role in phylogenetics. Sequence simulation has many applications, such as evaluating the performance of phylogenetic methods (Garland et al. 1993; Kuhner and Felsenstein 1994; Tateno et al. 1994; Huelsenbeck 1995), conducting parametric bootstraps, testing hypothesis (Goldman 1993a; Goldman 1993b; Adell and Dopazo 1994; Schoeniger and von Haeseler 1999), and generating data for training machine-learning applications (Abadi et al. 2020; Leuchtenberger et al. 2020; Ling et al. 2020; Suvorov et al. 2020). Typical sequence simulation programs (such as Seq-Gen (Rambaut and Grassly 1997), INDELible (Fletcher and Yang 2009), and Dawg (Cartwright 2005)) start with a given tree and a model of sequence evolution to generate an alignment of sequences at the tips of the tree (Figure 1A).

**Figure 1.**
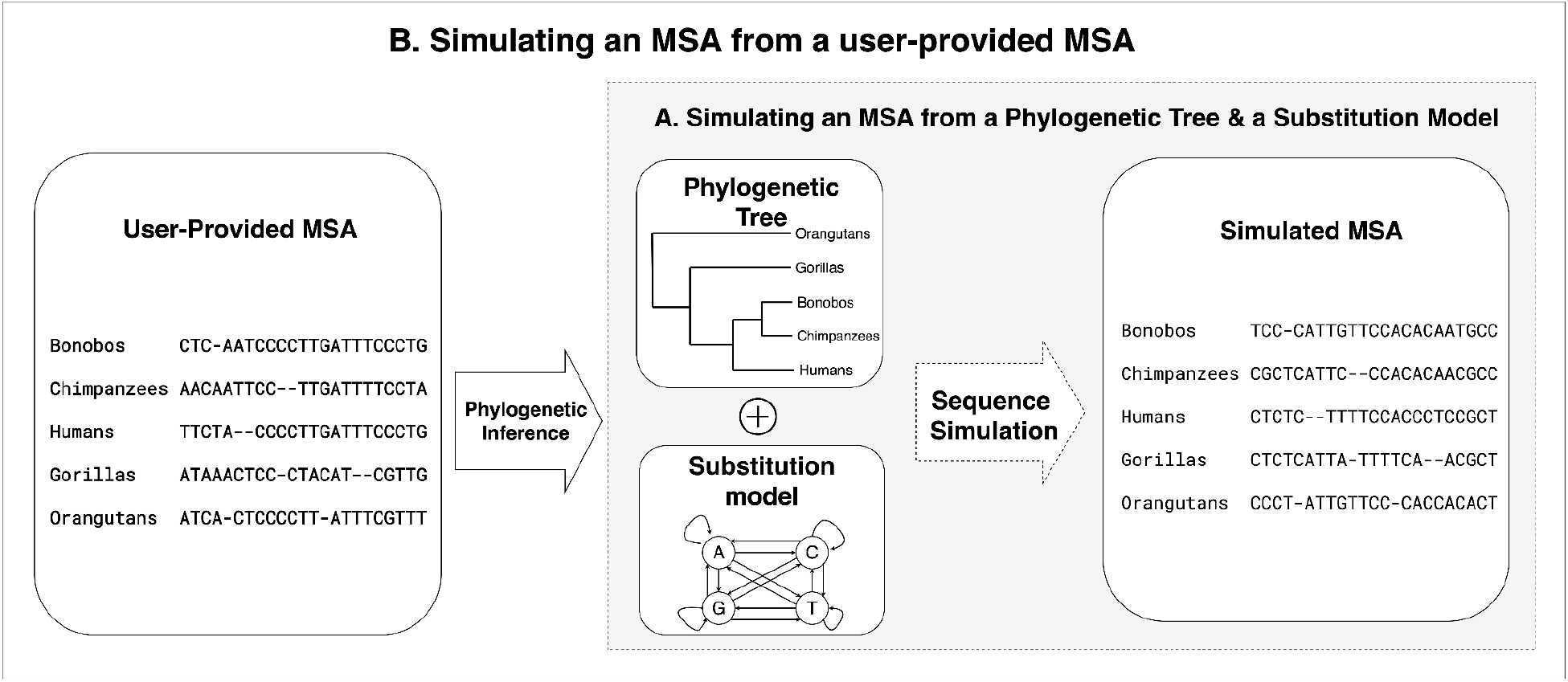
Sequence simulation process with two scenarios: (A) Simulating an MSA from a phylogenetic tree and a Markov substitution model, and (B) Simulating an MSA that mimics the underlying evolutionary process of a user-provided MSA. Here, the phylogenetic tree and the substitution model parameters are internally inferred from the user-provided MSA, which are used to simulate a new MSA.

Existing simulators often require long runtimes and a lot of memory to generate huge MSAs with millions of sequences or sites. Only phastSim (Maio et al. 2021), designed to simulate the alignments of genomes from viruses such as SARS-CoV-2, was recently introduced to efficiently simulate more than 100,000 viral sequences.

### New Approaches

Here, we develop a fast, efficient, versatile, and realistic sequence alignment simulator called AliSim. Our simulator integrates a wide range of evolutionary models, available in the IQ-TREE software (Nguyen et al. 2015; Minh et al. 2020), including standard, mixture, partition, and insertion-deletion models. Additionally, AliSim can simulate MSAs that mimic the evolutionary processes underlying empirical alignments, a feature not available in other tools. AliSim allows the user to provide an input MSA, then infers the evolutionary process from that MSA and subsequently simulates new MSAs from the inferred tree and model (Figure 1B). To further simplify this process, we also include the ability to simulate alignments based on the empirically-derived stationary distribution of nucleotides extracted from a large pre-computed database (Naser-Khdour et al. 2021). To reduce the runtime across a wide range of simulation conditions, we implement a new adaptive approach that allows AliSim to dynamically switch between the rate matrix approach (Schoeniger and von Haeseler 1995; Fletcher and Yang 2009) and the probability matrix approach (Schoeniger and von Haeseler 1995) during the simulation. As a result, AliSim can simulate large alignments with millions of sequences and sites using much lower computing times and memory than existing tools. For example, AliSim consumes 1.3 GB RAM and 1.3 hours to produce a COVID-19-like MSA containing one million sequences with thirty thousand sites per sequence. Whereas INDELible, Seq-Gen, and Dawg require two to five hours and 50 to 500 GB of RAM.

## Results

### AliSim supports a wide range of evolutionary models

Table 1 lists the full features of AliSim compared to other software. Notably, AliSim supports many evolutionary models not available in other software (Table 1). AliSim allows users to simulate different data types, including DNA, amino-acid, codon, binary, and multi-state morphological data using more than 200 time-reversible substitution models and 100 non-reversible models (Minh et al. 2020). AliSim also supports insertion-deletion models, as well as complex partition and mixture models. Moreover, users can specify model parameters or define new models via a short command-line option or a NEXUS file.

**Table 1.** Feature comparison between AliSim and existing tools, Seq-gen, INDELible, Dawg, and phastSim. (*) all partitions must share the same evolutionary model; (**) a mixture model is a set of substitution models where each site has a probability of belonging to a substitution model; (***) users can specify different evolutionary models to individual branches of a tree.

| Features | Seq-Gen | Dawg | INDELible | phastSim | AliSim |
| --- | --- | --- | --- | --- | --- |
| **Substitution models** |  |  |  |  |  |
| DNA | ✓ | ✓ | ✓ | ✓ | ✓ |
| Amino-acid | ✓ | ✓ | ✓ |  | ✓ |
| Codon |  | ✓ | ✓ | ✓ | ✓ |
| Binary and discrete morphological |  |  |  |  | ✓ |
| RNA (base-pairing) |  | ✓ |  |  |  |
| Non-reversible DNA and Amino-acid |  |  | ✓ | ✓ | ✓ |
| **Models of rate heterogeneity across sites** |  |  |  |  |  |
| Invariant sites (+I) | ✓ | ✓ | ✓ | ✓ | ✓ |
| Discrete Gamma distribution (+Gk) | ✓ |  | ✓ |  | ✓ |
| Continuous Gamma distribution (+GC) | ✓ | ✓ | ✓ | ✓ | ✓ |
| Distribution-free (+Rk) (user-defined) |  |  |  | ✓ | ✓ |
| Codon-position-specific rates | ✓ |  |  |  |  |
| Nonsynonymous/synonymous codon rate heterogeneity |  |  | ✓ | ✓ |  |
| **Complex models** |  |  |  |  |  |
| Insertion-deletion |  | ✓ | ✓ | ✓ | ✓ |
| Partition | Same model (*) | ✓ | ✓ |  | ✓ |
| Site mixture (**) |  |  | Codon only |  | ✓ |
| Tree mixture for non-treelike evolution | Same model and taxa | ✓ | ✓ |  | ✓ |
| Branch-specific substitutions (***) |  | ✓ | ✓ |  | ✓ |
| Hypermutability |  |  |  | ✓ |  |
| Heterotachy (Crotty et al. 2020) |  |  |  |  | ✓ |
| Functional divergence (Gaston et al. 2011) |  |  |  |  | ✓ |
| User-defined models | ✓ | ✓ | ✓ | ✓ | ✓ |
| Ascertainment Bias Correction |  |  |  |  | ✓ |
| **Biologically realistic simulations** |  |  |  |  |  |
| Mimicking a user-provided MSA |  |  |  |  | ✓ |
| Model parameters following empirical or user-defined distributions |  |  |  |  | ✓ |
| Simulating random trees |  |  | ✓ |  | ✓ |
| **Other features** |  |  |  |  |  |
| Multifurcating trees |  | ✓ | ✓ | ✓ | ✓ |
| Branch length scaling | ✓ | ✓ | ✓ | ✓ | ✓ |
| Graphical user interface | ✓ |  |  |  |  |
| Outputting ancestral sequences | ✓ | N/A | ✓ | ✓ | ✓ |
| Output format | PHYLIP,<br>NEXUS,<br>FASTA | PHYLIP,<br>FASTA,<br>NEXUS,<br>CLUSTAL,<br>POO | PHYLIP,<br>FASTA,<br>NEXUS | PHYLIP,<br>FASTA,<br>NEWICK,<br>MAT, Info | PHYLIP,<br>FASTA |
| Inserting output header | ✓ |  | ✓ |  |  |
| Output compression |  |  |  |  | Gzip |
| Programming language | C | C++ | C++ | Python | C++ |

To model rate heterogeneity across sites, AliSim offers invariant sites, discrete and continuous Gamma distributions (Yang 1994; Gu et al. 1995), distribution-free rate models (Yang 1995; Soubrier et al. 2012), and the GHOST model (Crotty et al. 2020). AliSim also implements branch-specific substitution models, which assign different models of sequence evolution to individual branches of a tree. To mimic more complex evolutionary patterns, such as incomplete lineage sorting or recombination, AliSim extends the partition model by allowing different tree topologies for each partition.

### AliSim offers more realistic simulations

#### Scenario 1: simulating MSAs that mimic a user-provided MSA (Figure 1B)

A common use-case for alignment simulation software is that users want to simulate an MSA that mimics the evolutionary history of a given MSA, for example, because this is needed for parametric bootstrap analysis. Until now, this required at least a two-step process whereby users first inferred the tree and model in one piece of software, then used these as input to the MSA simulation tool. The resulting MSA often failed to capture many characteristics of the original MSA, such as the position of gaps and the sitespecific evolutionary rates. AliSim improves this process by first running IQ-TREE to infer an evolutionary model and a tree from the input MSA, and then immediately generating any number of simulated MSAs from the inferred tree and model with the same gap patterns of the original MSA. For simulations under a mixture model, AliSim randomly assigns a model component of the mixture to each site according to the site posterior probability distribution of the mixture. For site-frequency mixture models, AliSim applies the posterior mean site frequencies (Wang et al. 2018). Similarly, AliSim employs the posterior mean site rates to better reflect the underlying evolutionary rate variation across sites. All these mechanisms help produce simulated MSAs that better reflect the relevant features of the original MSAs.

#### Scenario 2: simulating MSAs from a random tree and/or random parameters from empirical/user-defined distributions

When using Seq-Gen or Dawg, users need to provide as input a tree with branch lengths. To avoid this sometimes cumbersome step, AliSim allows users to generate a random tree under biologically plausible models such as the Birth-Death model (Kendall 1948) and the Yule-Harding model (Yule 1925; Harding 1971). For the Yule-Harding model, users only need to specify the number of leaves of the tree. For the birth-death model, users need to additionally provide the speciation and extinction rate. Branch lengths are randomly generated from an exponential distribution with a mean of 0.1 (by default) or from a user-defined distribution specified by a list of numbers.

Where users wish to simulate alignments that mimic empirical MSAs in the absence of a set of input alignments, AliSim can generate a stationary distribution of nucleotides from empirical distributions that were previously estimated from a large collection of empirical datasets (Naser-Khdour et al. 2021). Other parameters, such as substitution rates, nonsynonymous/synonymous rate ratios, transition, and transversion rates, can be drawn from user-defined lists of numbers, allowing AliSim to incorporate arbitrary distributions for all simulation parameters.

### AliSim automatically chooses a simulation method to minimize the runtime

Existing simulators typically employ either the rate matrix approach or the probability matrix approach to evolve sequences along a tree (see Methods). However, their performance varies with different sequence lengths (*L*) and branch lengths (*t*). Therefore, AliSim automatically switches between the rate matrix and probability matrix approaches to optimize the computing time. To determine when to use each approach, we compared the runtime of the rate matrix approach with the probability matrix approach on simulations using different combinations of *L* and *t* (see Methods).

The simulation results showed that the rate matrix approach is generally faster than the probability matrix approach when *L* * *t* < *2.226* and *L* * *t* < *17.307* for the discrete and continuous rate heterogeneity models, respectively. Therefore, the adaptive approach will apply the rate matrix approach for those branches that satisfy this inequality; otherwise, it will apply the probability matrix approach.

### AliSim is fast and efficient across a range of conditions

Figures 2A and 2B show that AliSim is faster than Seq-Gen, INDELible, Dawg, and phastSim for a range of common phylogenomic simulation conditions. Specifically, we simulated ‘deep’ data with 30K sites and 10K to 1M sequences, and we simulated ‘long’ data with 30K sequences and 10K to 1M sites (see Methods). We note before presenting these results that phastSim is designed specifically to simulate data along trees with a large proportion of extremely short branches. The trees in these simulations do not match these conditions, and so one might expect phastSim to perform poorly here. We include phastSim here for completeness, and present a comparison of phastSim and AliSim under the conditions for which phastSim was designed later.

**Figure 2.**
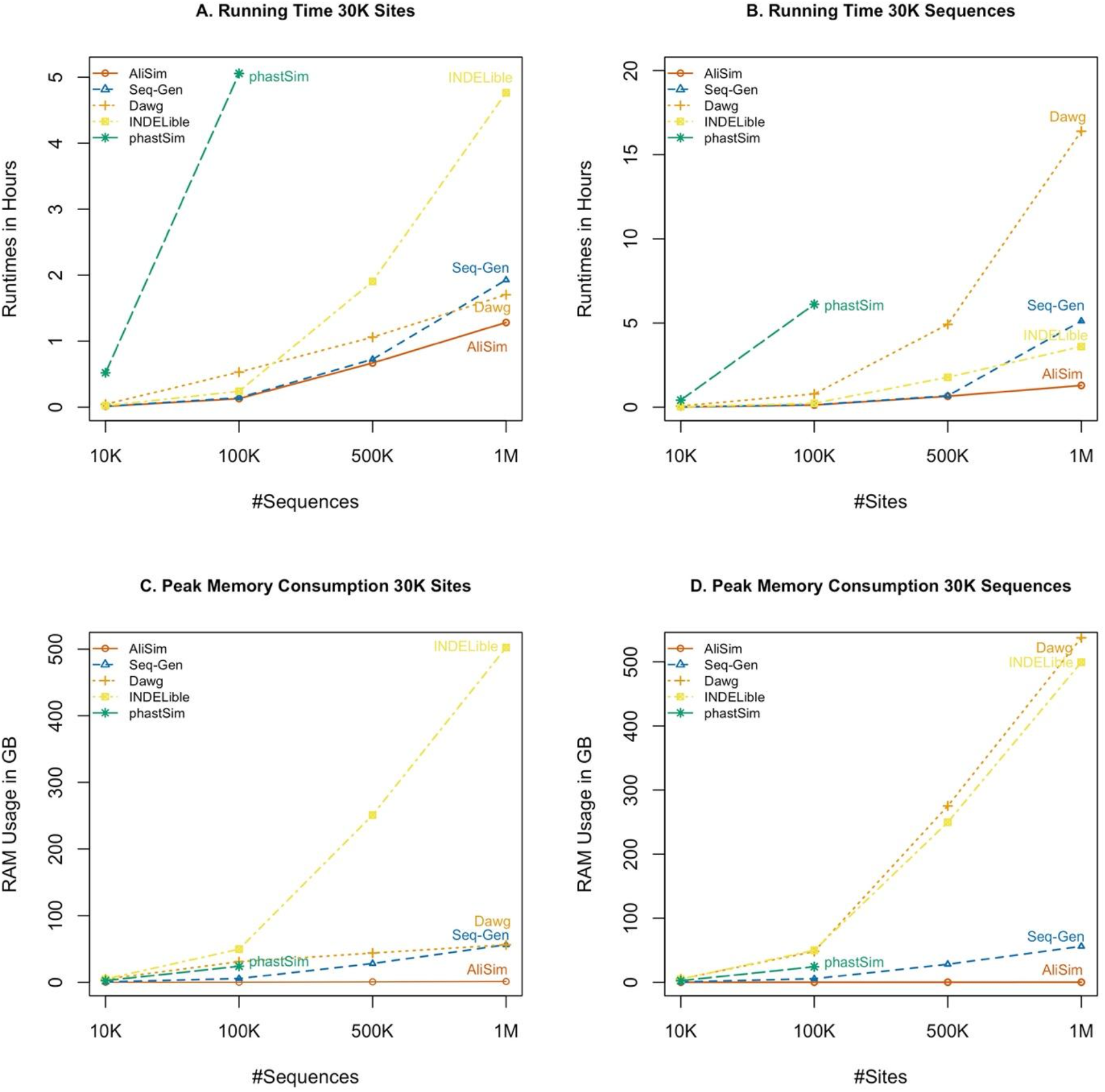
Benchmarking results of five software AliSim, Seq-Gen, Dawg, INDELible, and phastSim. Runtimes for (A) ‘deep’-data (varying number of sequences and 30K sites) and (B) ‘long’-data simulations (varying number of sites and 30K sequences) and peak memory consumption for (C) ‘deep’-data and (D) ‘long’-data simulations.

As expected, when the datasets were small, runtimes and memory usage were similar across all pieces of software. However, AliSim shows increasing advantages in runtimes and memory usage as the datasets get bigger. For example, for the deepest dataset (30K sites and 1M sequences), Seq-Gen, Dawg, INDELible, phastSim, and AliSim required 1.9, 1.7, 4.8, >24, and 1.3 hours of runtime, respectively. And for the longest dataset (30K sequences and 1M sites), Seq-Gen, Dawg, INDELible, phastSim, and AliSim required 5.1, 16.4, 3.6, >24, and 1.3 hours respectively. Thus, AliSim is a fast sequence simulator under a range of conditions.

AliSim shows dramatic improvements over other software in peak memory usage. Figures 2C and 2D show that these improvements become large even for fairly modestly-sized datasets. For the deepest dataset (30K sites and 1M sequences), Seq-Gen, Dawg, INDELible, and AliSim required 56.4, 56.3, 502.3, and 1.3 GB RAM, respectively (Figure 2C; phastSim peak memory usage was not recorded as it took >24hrs to run). For the longest dataset (30K sequences and 1M sites), Seq-Gen, Dawg, INDELible, and AliSim consumed 56, 537, 499, and 0.2 GB RAM, respectively (as above, phastSim was excluded because it took >24hrs to run). Importantly, the memory usage of AliSim only grows sub-linearly with respect to the dataset size. For the deep simulations, when increasing the number of sequences from 10K to 1M (100-fold), the RAM consumption of AliSim only increased from 137MB to 1.3GB (~10-fold increase, Figure 2C). For the long simulations, increasing the sequence length from 10K to 1M sites (100-fold) only increased the RAM usage marginally from 156MB to 222MB (less than a 2-fold increase, Figure 2D). This result is due to the memory saving technique in AliSim (see Methods), where the memory consumption is now dominated by storing the tree data structure and not by the simulated sequences.

We also tested the performance of these tools on simulating MSAs from COVID-19-like trees, which differ from commonly-simulated trees because they contain a large proportion of extremely short branch lengths, and for which phastSim was explicitly designed. In these conditions (Suppl. Figure S1A), phastSim and AliSim were the two fastest pieces of software, requiring 6 and 9 minutes respectively, whereas Seq-Gen, Dawg, and INDELible took 1.7, 1.7, and 3.2 hours respectively. In terms of RAM consumption, phastSim and AliSim only needed 1.4 and 1.3 GB RAM, respectively, while Seq-Gen, Dawg, and INDELible required 56, 51, and 502 GB RAM (Suppl. Figure S1B). We note that the performance of phastSim in these conditions is particularly remarkable because it is written in Python. Because of this, it may be that the language itself rather than the sequence simulation algorithms of phastSim are what limit its performance, and we speculate that phastSim may be able to be even faster if re-written in C or C++.

The adaptive approach helps AliSim achieve high performance by selecting the most efficient simulation approach for each branch. For example, in simulations under trees where branch lengths were generated from an exponential distribution with a mean of 0.1, the adaptive method will apply the probability matrix approach rather than the rate matrix approach on most branches, simply because most branches are longer than the switching parameters. The benefits of the adaptive approach can be measured in our simulations by forcing AliSim to use one method. For example, using the adaptive approach, AliSim took only 1.3 hours to simulate 1M sequences of 30K sites (Figure 2A). However, if we force AliSim to employ only the rate matrix approach, it takes more than 5 hours to simulate the same dataset. Similarly, the adaptive approach took only 9 minutes to simulate a dataset on a COVID-19-like tree (Suppl. Figure S1A), but if we force AliSim to use the probability matrix approach, the same simulation takes 1.3 hours.

### Software validation

To validate the AliSim, we simulated 287 MSAs with 100 sequences across a wide range of substitution models and insertion-deletion rates of 0.0, 0.02, 0.04, 0.06, 0.08 and 0.1. These choices of indel rates follow empirical studies (Cartwright 2009). We then ran IQ-TREE to determine the best-fit model using ModelFinder (Kalyaanamoorthy et al. 2017) and reconstructed phylogenetic trees under the best-fit model. We compared the topology between the true trees and the inferred trees using the Robinson-Foulds distance (Robinson and Foulds 1981).

Supplementary Table S1 showed that in 147 tests (51.22%), the true model was recovered as the best-fit model. In 243 tests (84.67%), 246 tests (85.71%), and 267 tests (93.03%), the true models appear in the top-2, top-3, and top-4 best models, respectively. The average Robinson-Foulds distance between the true trees and the inferred trees across all test cases was 1.99 (s.e. 0.133). That means the inferred trees differed from the true trees by only 0.995 out of 97 (1.03%) internal branches. The tree lengths (sum of branch lengths) of the inferred trees differed from the true trees by only 1.9%.

For simulations with non-zero indel rates, the average differences in the alignment length and proportion of gaps between MSAs simulated by AliSim and those by INDELible were 0.5% and 0.22%, respectively (Suppl. Table S2).

### Conclusion

In conclusion, AliSim is a fast and memory-efficient simulation tool, which simplifies and speeds up many common workflows in phylogenetics. Because AliSim has a small memory footprint, it can be used to simulate even very large alignments on personal computers.

## Materials and Methods

We developed AliSim in C++ as an extension to the IQ-TREE software to take advantage of all models of sequence evolution provided in IQ-TREE. Generally, AliSim works by first generating a sequence at the root of the tree following the stationarity of the model. AliSim then recursively traverses along the tree to generate sequences at each node of the tree based on the sequence of its ancestral node. AliSim completes this process once all the sequences at the tips are generated. In the following, we introduce general notations and three simulation approaches to simulate sequence evolution along a branch of a tree in a general case with insertions and deletions (indels).

Let *Q* = (*q_xy_*) be a rate matrix of a Markov model, where *x*, *y* ∈ *Σ*, a finite alphabet, for example, the alphabet of nucleotides or amino acids; *q_xy_* is the instantaneous substitution rate from *x* to *y*. *Q* is normalized such that the row sum is zero: *q_xx_* = –∑_*y*≠*x*_*q_xy_* and the total substitution rate is one: ∑_*x*_ *π_x_ q_xx_* = –*1*, where *π_x_* is the state frequency. We assume a model of rate heterogeneity across sites, such as the invariant site proportion, the continuous/discrete Gamma model (Yang 1994), or the distribution-free rate model (Yang 1995; Soubrier et al. 2012). Let *r_I_*, *r_D_* be the insertion and deletion rate, respectively, relative to the substitution rate. Let *Φ_I_*, *Φ_D_* be the insertion-size and deletion-size distributions, respectively. AliSim allows users to use built-in indel-size distributions, such as Geometric, Negative Binomial, Zipfian, and Lavalette distribution (Fletcher and Yang 2009), or specify their own distributions. By default, AliSim uses a Zipfian distribution with an exponent of 1.7 as previously estimated from empirical data (Steven A.Benner et al. 1993; Cartwright 2009). Given a sequence *X* = *x_1_x_2_*…*x_L_*,*x_i_* ∈ *Σ* ∪ {–} (where ′ – ′ denotes the gap character), at an ancestral node of a phylogenetic tree; a vector of site-specific rate *R* = *r_1_r_2_*… *r_L_*, *r_i_* is generated according to the site-rate heterogeneity model; and a branch length *t*, as the number of substitutions per site, of a branch connecting the ancestral node to a descendent node, we now describe three approaches to generate a new sequence *Y* = *y_1_y_2_*… *y_L′_*, *y_i_* ∈ ∑ ∪ {–}, at the descendent node. *L*′ might be different from *L* if the insertion rate is non-zero.

### The rate matrix approach

This approach implements the Gillespie algorithm (Gillespie 1977) as follows. We compute the total mutation rate for the ancestral sequence *X* as the sum of site-specific mutation rates: *M* = *S* + *I* + *D*, where *S*, *I*, *D* is the total rate of substitutions, insertions, and deletions of all sites respectively. 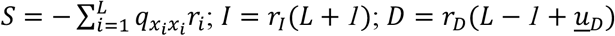, where *<u>u</u>_D_* is the mean of the deletion size distribution (Cartwright 2005).

0. Set *Y* ← *X* and *L*′ ← *L*.
1. Generating a waiting time *w* for a mutation to occur from an exponential distribution with a mean of 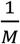.
2. If *w* > *t*, no mutation occurs, then we stop and return *Y* as the sequence at the descendent node.
3. If *w* ≤ *t*, a mutation occurs and we randomly select a mutation type as substitution, insertion, or deletion with probabilities of 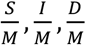 respectively.
4. If the mutation type is substitution:

4.1. Randomly select a non-gap position *i*, *1* ≤ *i*. ≤ *L*′ where the substitution occurs with probabilities 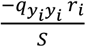. If *Y* contains only gaps, we terminate the algorithm.
4.2. Randomly choose a new state *z_i_* (according to probabilities 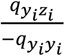.
4.3. Update the total substitution rate to reflect the new state: *S* ← *S* + (*q_y_i_y_i__* – *q_z_i_z_i__*)^r_i_^.
4.4. Assign *y_i_* ← *z_i_* and go to step 7.
5. If the mutation type is insertion:

5.1. Uniformly select a non-gap position *i*, *1* ≤ *i* ≤ *L*′ + *1* where the insertion occurs.
5.2. Randomly generate a new sequence *Z* = *Z_1_*…*Z_j_* based on the stationary distribution of the model, where the sequence length *j* follows the insertion-size distribution *Φ_I_*.
5.3. Insert *Z* into *Y* at position *i*. If *i* = *L*′ + *1*, *Z* is appended at the end of *Y*.
5.4. Insert a stretch of *j* gaps into position *i*. of the sequences at all other nodes of the tree so that all sequences have the same length.
5.5. Generate a vector of site rates (*s_1_*,…, *S_j_*) according to the distribution of rate heterogeneity across sites and insert this vector into *R*, at position *i*.
5.6. Update the sequence length *L*′ ← *L*′ + *j*, the total substitution rate 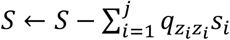, the total insertion rate *I* ← *I* + *r_I_j*, and the total deletion rate *D* ← *D* + *r_D_j*.
5.7. Go to step 7.
6. If the mutation type is deletion:

6.1. Generate a deletion length, *j*, from the deletion-size distribution *Φ_D_*.
6.1. Uniformly select a non-gap position *i*, *1* ≤ *i* ≤ *L*′ – *j* + *1*, where the deletion occurs.
6.3. Initialize *P* = {*p_1_*, *p_2_*,…,*p_j_*}, a set of *j* non-gap positions in *Y* starting at position *i*. Note that *p_j_* might be greater than *i* + *j* if there are gaps between position *i* and *i* + *j*.
6.4. Update the total substitution rate 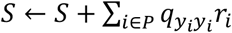; then ∀*i* ∈ *P*, we set *y_i_* ← ′ – ′ and *r_i_* ← *0*.
6.5. Update the total insertions rate *I* ← *I* – *r_I_j*, and the deletion rate *D* ← *D* – *r_D_j*.
7. Update the total mutation rate: *M* ← *S* + *I* + *D* and the time *t* ← *t* – *w*. Go back to step 1.

### The probability matrix approach

Instead of generating a series of waiting times, the probability matrix approach generates a new state *y_i_* for each site in the sequence based on the state *x_i_*, *i* = *1*,*2*,…, *L*. For each site *i*, we compute the transition probability matrix *P*(*t*, *r_i_*) = *e^Qtr_i_^*. Then, the new state *y_i_* is drawn from the probability distribution *P_x_i_y_i__*(*t*, *r_i_*), *y_i_* ∈ *Σ*. Note that when using a discrete rate model with *k* categories, we only need to compute *P*(*t*, *r_i_*) exactly *k* times to save computations. Whereas for a continuous Gamma rate model, we have to compute *P*(*t*, *r_i_*) for each site independently. After processing substitutions with the probability matrix approach, to simulate indels, we apply the Gillespie algorithm as described above on the new sequence *Y* without considering substitutions by setting and maintaining the total substitution rate *S* at zero.

### The adaptive approach

In simulations without indels, the probability matrix approach has a time complexity independent of branch lengths, but the time complexity for the rate matrix approach grows with increasing branch lengths. In simulations with indels, the branch lengths affect the runtime of the rate matrix approach more significantly than that of the probability matrix approach. We expect the rate matrix approach to outperform the probability matrix approach for small *t* but the opposite for large *t*. Therefore, we derived an adaptive approach, in which we determined a switching parameter from the sequence length. For all branches where the branch length is smaller than this parameter, we employ the rate matrix approach. For the remaining (long) branches, we use the probability matrix approach. That means our adaptive algorithm will automatically switch between these two approaches on a per-branch basis to minimize the computations.

To determine the switching parameter, we performed simulations with different sequence lengths, ranging from 1K to 100K sites (a total of 19 tests), with/without rate heterogeneity. We measured the runtimes of the probability matrix approach when simulating MSAs under a random Yule-Harding tree with 10K tips based on the general time-reversible (GTR) model (Tavaré 1986) with/without continuous Gamma rate heterogeneity using a Gamma shape of 0.5. For each test case, we applied binary search on a predefined range of branch length to determine the switching parameter where the runtime of the rate matrix approach is less than the probability matrix approach (Suppl. Table S3). We then determine the switching parameters using a least square fit across the simulations.

### Memory optimization techniques

Naively, when simulating sequences along a bifurcating tree with *n* tips, we need to store up to (2*n*-1) sequences, consisting of (*n* – 1) internal nodes and *n* tips. To save memory, we release the memory allocated to the sequence of an internal node if the sequences of its children nodes are already generated. Additionally, in simulations without partitions and indels, AliSim writes out the tip sequences to the output file immediately after simulating them, then frees the memory. This approach considerably reduces the maximum number of sequences that need to be stored in memory from (2*n* – 1) to the maximum depth of the tree. For a balanced bifurcating tree, this maximal depth is *log*_2_(*n*) + 1, leading to a substantial reduction in memory usage.

### Benchmark experimental setup

We benchmarked AliSim against Seq-Gen, INDELible, Dawg, and phastSim. For INDELible, we used Method 2 which is more efficient than Method 1 in simulations with continuous rate heterogeneity across sites (Fletcher and Yang 2009). The benchmark was run on a Linux server with 2.0 GHz AMD EPYC 7501 32-Core Processor and 1-TB RAM. Inspired by the newly emerged COVID-19 data, we tested the ability of all tools to simulate alignments with 30K sites and 10K, 100K, 500K, and 1M DNA sequences. For the model of sequence evolution, we applied the GTR model with a 0.2 proportion of invariant sites and a continuous Gamma model of rate heterogeneity across sites (shape parameter of 0.5). We ran different software to simulate sequences along a random Yule-Harding tree with exponentially distributed branch lengths with a mean of 0.1. We call this the ‘deep’-data simulation. Moreover, to mimic the size of real phylogenomic datasets, we simulated MSAs with 30K sequences and increased sequence length from 10K to 1M sites. This is called ‘long’-data simulation.

## Supplementary Material

Supplementary data are available at Molecular Biology and Evolution online. Simulated data, testing scripts, and benchmark results in this article are all available at https://doi.org/10.5281/zenodo.5778560.

## Acknowledgments

This work was supported by a Chan-Zuckerberg Initiative grant for open source software for science to B.Q.M. and R.L., an Australian Research Council Discovery Grant [DP200103151 to R.L. and B.Q.M.], and partly by a Vingroup Science and Technology Scholarship [VGRS20042M to N.T.L.]. The computational results have been obtained on the cluster at the Center for Integrative Bioinformatics Vienna (CIBIV). We thank Arndt von Haeseler for valuable comments and providing access to the CIBIV cluster; Caitlin Cherryh, Yu Lin, Fred Jaya, Weiwen Wang for their comments on the manuscript; Andrew Roger and Edward Susko for helpful discussions.

## References

Abadi S, Avram O, Rosset S, Pupko T, Mayrose I. 2020. ModelTeller: Model selection for optimal phylogenetic reconstruction using machine learning. Mol Biol Evol. 37(11):3338–3352.

Adell JC, Dopazo J. 1994. Monte Carlo simulation in phylogenies: An application to test the constancy of evolutionary rates. J Mol Evol. 38(3):305–309.

Cartwright RA. 2005. DNA assembly with gaps (Dawg): Simulating sequence evolution. Bioinformatics. 21(SUPPL. 3):31–38.

Cartwright RA. 2009. Problems and solutions for estimating indel rates and length distributions. Mol Biol Evol. 26(2):473–480.

Crotty SM, Minh BQ, Bean NG, Holland BR, Tuke J, Jermiin LS, von Haeseler A. 2020. GHOST: Recovering Historical Signal from Heterotachously Evolved Sequence Alignments. Syst Biol. 69(2):249–264.

De Maio Nicola, Weilguny L, Walker CR, Turakhia Y, Corbett-Detig R, Goldman N. 2021. phastSim: efficient simulation of sequence evolution for pandemic-scale datasets. bioRxiv. doi: 10.1101/2021.03.15.435416.

Fletcher W, Yang Z. 2009. INDELible: A flexible simulator of biological sequence evolution. Mol Biol Evol. 26(8):1879–1888.

Garland T, Dickerman AW, Janis CM, Jones JA. 1993. Phylogenetic Analysis of Covariance by Computer Simulation. Syst Biol. 42(3):265–292.

Gaston D, Susko E, Roger AJ. 2011. A phylogenetic mixture model for the identification of functionally divergent protein residues. Bioinformatics. 27(19):2655–2663.

Gillespie DT. 1977. Exact stochastic simulation of coupled chemical reactions. J Phys Chem. 81(25):2340–2361.

Goldman N. 1993a. Statistical tests of models of DNA substitution. J Mol Evol. 36(2):182–198.

Goldman N. 1993b. Simple diagnostic statistical tests of models for DNA substitution. J Mol Evol. 37(6):650–661.

Gu X, Fu Y-X, Li W-H. 1995. Maximum likelihood estimation of the heterogeneity of substitution rate among nucleotide sites. Mol Biol Evol. 2(4):546–557.

Harding EF. 1971. The Probabilities of Rooted Tree-Shapes Generated by Random Bifurcation. Adv Appl Probab. 3(1):44–77.

Huelsenbeck JP. 1995. Performance of Phylogenetic Methods in Simulation. Syst Biol. 44(1):17–48.

Kalyaanamoorthy S, Minh BQ, Wong TKF, von Haeseler A, Jermiin LS. 2017. ModelFinder: Fast model selection for accurate phylogenetic estimates. Nat Methods. 14(6):587–589.

Kendall DG. 1948. On the Generalized “Birth-and-Death” Process. Ann Math Stat. 19(1):1–15.

Kuhner MK, Felsenstein J. 1994. A simulation comparison of phylogeny algorithms under equal and unequal evolutionary rates. Mol Biol Evol. 11(3):459–468.

Leuchtenberger AF, Crotty SM, Drucks T, Schmidt HA, Burgstaller-Muehlbacher S, von Haeseler A. 2020. Distinguishing Felsenstein Zone from Farris Zone Using Neural Networks. Mol Biol Evol. 37(12):3632–3641.

Ling C, Cheng W, Zhang Haoyu, Zhu H, Zhang Hua. 2020. Deep Neighbor Information Learning from Evolution Trees for Phylogenetic Likelihood Estimates. IEEE Access. 8:220692–220702.

Minh BQ, Schmidt HA, Chernomor O, Schrempf D, Woodhams MD, von Haeseler A, Lanfear R. 2020. IQ-TREE 2: New Models and Efficient Methods for Phylogenetic Inference in the Genomic Era. Mol Biol Evol. 37(5):1530–1534.

Naser-Khdour S, Minh BQ, Robert L. 2021. The Influence of Model Violation on Phylogenetic Inference: A Simulation Study. bioRxiv.

Nguyen LT, Schmidt HA, von Haeseler A, Minh BQ. 2015. IQ-TREE: A fast and effective stochastic algorithm for estimating maximum-likelihood phylogenies. Mol Biol Evol. 32(1):268–274.

Rambaut A, Grassly NC. 1997. Seq-gen: An application for the monte carlo simulation of dna sequence evolution along phylogenetic trees. Bioinformatics. 13(3):235–238.

Robinson DF, Foulds LR. 1981. Comparison of phylogenetic trees. Math Biosci. 53(1–2):131–147.

Schoeniger M, von Haeseler A. 1995. Simulating efficiently the evolution of DNA sequences. Bioinformatics. 11(1):111–115.

Schoeniger M, von Haeseler A. 1999. Toward Assigning Helical Regions in Alignments of Ribosomal RNA and Testing the Appropriateness of Evolutionary Models. J Mol Evol Vol. 49:691–698.

Soubrier J, Steel M, Lee MSY, Der Sarkissian C, Guindon S, Ho SYW, Cooper A. 2012. The influence of rate heterogeneity among sites on the time dependence of molecular rates. Mol Biol Evol. 29(11):3345–3358.

Steven A. Benner, Mark A. Cohen, Gaston H. Gonnet. 1993. Empirical and Structural Models for Insertions and Deletions in the Divergent Evolution of Proteins. J Mol Biol. 229(4):1065–1082.

Suvorov A, Hochuli J, Schrider DR. 2020. Accurate Inference of Tree Topologies from Multiple Sequence Alignments Using Deep Learning. Syst Biol. 69(2):221–233.

Tateno Y, Takezaki N, Nei M. 1994. Relative efficiencies of the maximum-likelihood, neighbor-joining, and maximum-parsimony methods when substitution rate varies with site. Mol Biol Evol. 11(2):261–277.

Tavaré S. Miura RM 1986. Some probabilistic and statistical problems in the analysis of DNA sequences. Lect Math life Sci. 17:57–86.

Wang HC, Minh BQ, Susko E, Roger AJ. 2018. Modeling Site Heterogeneity with Posterior Mean Site Frequency Profiles Accelerates Accurate Phylogenomic Estimation. Syst Biol. 67(2):216–235.

Yang Z. 1994. Maximum likelihood phylogenetic estimation from DNA sequences with variable rates over sites: Approximate methods. J Mol Evol. 39(3):306–314.

Yang Z. 1995. A space-time process model for the evolution of DNA sequences. Genetics. 139(2):993–1005.

Yule GU. 1925. A Mathematical Theory of Evolution Based on the Conclusions of Dr. J. C. Willis, F.R.S. Philos Trans R Soc London Ser B, Contain Pap a Biol Character. 213:21–87.

