## Supplementary materials for "AliSim: A Fast and Versatile Phylogenetic Sequence Simulator For the Genomic Era"

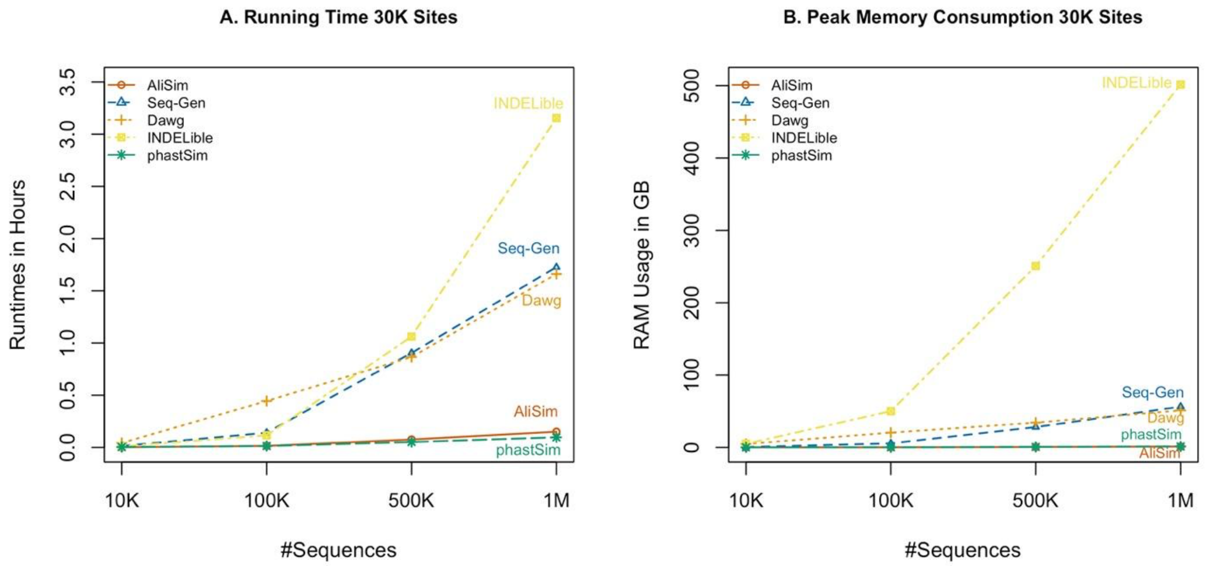


**Figure S1.** Benchmarking results of five software AliSim, Seq-Gen, Dawg, phastSim, and INDELible in COVID-like trees with extremely short branch lengths: Runtimes (A) and peak memory consumption (B). Here, the input trees are drawn under a Yule process with the birth rate equal to one divided by the output sequence length so that there is on average only one mutation per branch (De Maio et al. 2021).

**Table S1.** Validation results of AliSim on 287 simulations across a wide range of substitution models and insertion-deletion rates of 0.0, 0.02, 0.04, 0.06, 0.08 and 0.1.

| No | Data Type | Indel-rate | The true (input) model | The best-fit model | Rank of the true model | RF distance | Inferred tree length | True tree length |
|---|---|---|---|---|---|---|---|---|
| Substitution models | | | | | | | | |
| 1 | DNA | 0/0 | GTR{2.0,36.5,50.3,10.0,20.7}+F{0.3,0.1,0.2,0.4} | GTR+F+ASC | 2 | 4 | 21.278 | 21.385 |
| 2 | DNA | 0/0 | GTR{2.0,36.5,50.3,10.0,20.7}+F{0.2,0.1,0.3,0.4} +I{0.2} | GTR+F+I | 1 | 0 | 21.101 | 19.890 |
| 3 | DNA | 0/0 | GTR{2.0,36.5,50.3,10.0,20.7}+F{0.4,0.1,0.2,0.3} +G{0.5} | GTR+F+G4 | 1 | 0 | 19.654 | 19.769 |
| 4 | DNA | 0/0 | GTR{2.0,36.5,50.3,10.0,20.7}+F{0.1,0.2,0.4,0.3} +I{0.2}+G{0.5} | GTR+F+I+G4 | 1 | 2 | 20.309 | 20.194 |
| 5 | DNA | 0/0 | JC | JC | 1 | 4 | 19.733 | 19.706 |
| 6 | DNA | 0/0 | JC+I{0.2} | JC+I | 1 | 2 | 19.695 | 21.315 |
| 7 | DNA | 0/0 | JC+G{0.5} | JC+G4 | 1 | 6 | 21.483 | 21.543 |
| 8 | DNA | 0/0 | JC+I{0.2}+G{0.5} | JC+I+G4 | 1 | 0 | 18.669 | 18.706 |
| 9 | DNA | 0/0 | F81+F{0.3,0.1,0.2,0.4} | F81+F+ASC | 2 | 4 | 20.174 | 20.150 |
| 10 | DNA | 0/0 | F81+F{0.2,0.1,0.3,0.4}+I{0.2} | F81+F+I | 1 | 2 | 19.627 | 19.341 |
| 11 | DNA | 0/0 | F81+F{0.4,0.1,0.2,0.3}+G{0.5} | F81+F+G4 | 1 | 0 | 19.646 | 19.811 |
| 12 | DNA | 0/0 | F81+F{0.1,0.2,0.4,0.3}+I{0.2}+G{0.5} | F81+F+I+G4 | 1 | 2 | 18.882 | 18.877 |
| 13 | DNA | 0/0 | K80{2.0} | K2P | 1 | 0 | 21.212 | 21.255 |
| 14 | DNA | 0/0 | K80{2.0}+I{0.2} | K2P+I | 1 | 2 | 16.908 | 17.401 |
| 15 | DNA | 0/0 | K80{2.0}+G{0.5} | K2P+G4 | 1 | 0 | 19.947 | 19.986 |
| 16 | DNA | 0/0 | K80{2.0}+I{0.2}+G{0.5} | K2P+I+G4 | 1 | 6 | 20.690 | 20.546 |
| 17 | DNA | 0/0 | HKY{2.0}+F{0.3,0.1,0.2,0.4} | HKY+F+ASC | 2 | 2 | 22.092 | 22.037 |

| | | | | | | | | |
|---|---|---|---|---|---|---|---|---|
| 18 | DNA | 0/0 | HKY{2.0}+F{0.2,0.1,0.3,0.4}+I{0.2} | HKY+F+I | 1 | 2 | 19.521 | 21.586 |
| 19 | DNA | 0/0 | HKY{2.0}+F{0.4,0.1,0.2,0.3}+G{0.5} | HKY+F+G4 | 1 | 0 | 17.866 | 18.032 |
| 20 | DNA | 0/0 | HKY{2.0}+F{0.1,0.2,0.4,0.3}+I{0.2}+G{0.5} | TPM2u+F+I+G4 | 2 | 2 | 20.959 | 21.021 |
| 21 | DNA | 0/0 | TN{2.0,4.0}+F{0.3,0.1,0.2,0.4} | TN+F+ASC | 2 | 0 | 20.641 | 20.676 |
| 22 | DNA | 0/0 | TN{2.0,4.0}+F{0.2,0.1,0.3,0.4}+I{0.2} | TN+F+I | 1 | 2 | 20.969 | 20.645 |
| 23 | DNA | 0/0 | TN{2.0,4.0}+F{0.4,0.1,0.2,0.3}+G{0.5} | TN+F+G4 | 1 | 4 | 19.989 | 20.025 |
| 24 | DNA | 0/0 | TN{2.0,4.0}+F{0.1,0.2,0.4,0.3}+I{0.2}+G{0.5} | TN+F+I+G4 | 1 | 4 | 21.945 | 21.939 |
| 25 | DNA | 0/0 | TNe{2.0,4.0} | TNe+ASC | 2 | 0 | 20.485 | 20.511 |
| 26 | DNA | 0/0 | TNe{2.0,4.0}+I{0.2} | TNe+I | 1 | 4 | 15.721 | 19.082 |
| 27 | DNA | 0/0 | TNe{2.0,4.0}+G{0.5} | TNe+G4 | 1 | 0 | 19.682 | 18.843 |
| 28 | DNA | 0/0 | TNe{2.0,4.0}+I{0.2}+G{0.5} | TNe+I+G4 | 1 | 4 | 21.828 | 21.728 |
| 29 | DNA | 0/0 | K81{2.0,4.0} | K3P+ASC | 2 | 0 | 20.321 | 20.325 |
| 30 | DNA | 0/0 | K81{2.0,4.0}+I{0.2} | K3P+I | 1 | 0 | 16.045 | 20.755 |
| 31 | DNA | 0/0 | K81{2.0,4.0}+G{0.5} | K3P+G4 | 1 | 2 | 18.938 | 18.966 |
| 32 | DNA | 0/0 | K81{2.0,4.0}+I{0.2}+G{0.5} | K3P+I+G4 | 1 | 4 | 21.038 | 21.135 |
| 33 | DNA | 0/0 | K81u{2.0,4.0}+F{0.3,0.1,0.2,0.4} | K3Pu+F+ASC | 2 | 0 | 18.344 | 18.450 |
| 34 | DNA | 0/0 | K81u{2.0,4.0}+F{0.2,0.1,0.3,0.4}+I{0.2} | K3Pu+F+I | 1 | 6 | 20.120 | 21.044 |
| 35 | DNA | 0/0 | K81u{2.0,4.0}+F{0.4,0.1,0.2,0.3}+G{0.5} | K3Pu+F+G4 | 1 | 0 | 18.970 | 19.023 |
| 36 | DNA | 0/0 | K81u{2.0,4.0}+F{0.1,0.2,0.4,0.3}+I{0.2}+G{0.5} | K3Pu+F+I+G4 | 1 | 0 | 20.168 | 20.138 |
| 37 | DNA | 0/0 | TPM2{2.0,4.0} | TPM2+ASC | 2 | 0 | 19.333 | 19.401 |
| 38 | DNA | 0/0 | TPM2{2.0,4.0}+I{0.2} | TPM2+I | 1 | 0 | 19.165 | 20.671 |
| 39 | DNA | 0/0 | TPM2{2.0,4.0}+G{0.5} | TPM2+G4 | 1 | 2 | 19.309 | 19.355 |
| 40 | DNA | 0/0 | TPM2{2.0,4.0}+I{0.2}+G{0.5} | TPM2+I+G4 | 1 | 0 | 20.994 | 20.874 |
| 41 | DNA | 0/0 | TPM2u{2.0,4.0}+F{0.3,0.1,0.2,0.4} | TPM2u+F+ASC | 2 | 0 | 18.087 | 18.118 |

| | | | | | | | |
|---|---|---|---|---|---|---|---|
| 42 DNA | 0/0 | TPM2u{2.0,4.0}+F{0.2,0.1,0.3,0.4}+I{0.2} | TPM2u+F+I | 1 | 2 | 19.714 | 21.628 |
| 43 DNA | 0/0 | TPM2u{2.0,4.0}+F{0.4,0.1,0.2,0.3}+G{0.5} | TPM2u+F+G4 | 1 | 0 | 19.905 | 19.956 |
| 44 DNA | 0/0 | TPM2u{2.0,4.0}+F{0.1,0.2,0.4,0.3}+I{0.2}+G{0.5} | TPM2u+F+I+G4 | 1 | 6 | 21.678 | 21.745 |
| 45 DNA | 0/0 | TPM3{2.0,4.0} | TPM3+ASC | 2 | 0 | 19.080 | 19.057 |
| 46 DNA | 0/0 | TPM3{2.0,4.0}+I{0.2} | TPM3+I | 1 | 4 | 18.682 | 20.628 |
| 47 DNA | 0/0 | TPM3{2.0,4.0}+G{0.5} | TPM3+G4 | 1 | 4 | 20.349 | 20.445 |
| 48 DNA | 0/0 | TPM3{2.0,4.0}+I{0.2}+G{0.5} | TPM3+I+G4 | 1 | 4 | 20.238 | 20.227 |
| 49 DNA | 0/0 | TPM3u{2.0,4.0}+F{0.3,0.1,0.2,0.4} | TPM3u+F+ASC | 2 | 2 | 20.047 | 19.996 |
| 50 DNA | 0/0 | TPM3u{2.0,4.0}+F{0.2,0.1,0.3,0.4}+I{0.2} | TPM3u+F+I | 1 | 0 | 21.102 | 19.425 |
| 51 DNA | 0/0 | TPM3u{2.0,4.0}+F{0.4,0.1,0.2,0.3}+G{0.5} | TPM3u+F+G4 | 1 | 2 | 18.297 | 18.307 |
| 52 DNA | 0/0 | TPM3u{2.0,4.0}+F{0.1,0.2,0.4,0.3}+I{0.2}+G{0.5} | TPM3u+F+I+G4 | 1 | 0 | 20.356 | 20.499 |
| 53 DNA | 0/0 | TIM{2.0,4.0,6.0}+F{0.3,0.1,0.2,0.4} | TIM+F+ASC | 2 | 6 | 20.878 | 20.975 |
| 54 DNA | 0/0 | TIM{2.0,4.0,6.0}+F{0.2,0.1,0.3,0.4}+I{0.2} | TIM+F+I | 1 | 10 | 17.254 | 18.367 |
| 55 DNA | 0/0 | TIM{2.0,4.0,6.0}+F{0.4,0.1,0.2,0.3}+G{0.5} | TIM+F+G4 | 1 | 0 | 19.885 | 19.890 |
| 56 DNA | 0/0 | TIM{2.0,4.0,6.0}+F{0.1,0.2,0.4,0.3}+I{0.2}+G{0.5} | TIM+F+I+G4 | 1 | 0 | 18.382 | 18.565 |
| 57 DNA | 0/0 | TIMe{2.0,4.0,6.0} | TIMe+ASC | 2 | 0 | 19.395 | 19.344 |
| 58 DNA | 0/0 | TIMe{2.0,4.0,6.0}+I{0.2} | TIMe+I | 1 | 4 | 19.829 | 18.809 |
| 59 DNA | 0/0 | TIMe{2.0,4.0,6.0}+G{0.5} | TIMe+G4 | 1 | 6 | 19.586 | 19.644 |
| 60 DNA | 0/0 | TIMe{2.0,4.0,6.0}+I{0.2}+G{0.5} | TIMe+I+G4 | 1 | 2 | 20.502 | 20.569 |
| 61 DNA | 0/0 | TIM2{2.0,4.0,6.0}+F{0.3,0.1,0.2,0.4} | TIM2+F+ASC | 2 | 4 | 17.652 | 17.697 |
| 62 DNA | 0/0 | TIM2{2.0,4.0,6.0}+F{0.2,0.1,0.3,0.4}+I{0.2} | TIM2+F+I | 1 | 6 | 20.355 | 21.957 |
| 63 DNA | 0/0 | TIM2{2.0,4.0,6.0}+F{0.4,0.1,0.2,0.3}+G{0.5} | TIM2+F+G4 | 1 | 0 | 18.273 | 18.273 |

| | | | | | | | | |
|---|---|---|---|---|---|---|---|---|
| 64 | DNA | 0/0 | TIM2{2.0,4.0,6.0}+F{0.1,0.2,0.4,0.3}+I{0.2}+G{0.5} | TIM2+F+I+G4 | 1 | 2 | 19.363 | 19.317 |
| 65 | DNA | 0/0 | TIM2e{2.0,4.0,6.0} | TIM2e+ASC | 2 | 8 | 18.393 | 18.371 |
| 66 | DNA | 0/0 | TIM2e{2.0,4.0,6.0}+I{0.2} | TIM2e+I | 1 | 2 | 15.628 | 19.874 |
| 67 | DNA | 0/0 | TIM2e{2.0,4.0,6.0}+G{0.5} | TIM2e+G4 | 1 | 2 | 19.427 | 19.529 |
| 68 | DNA | 0/0 | TIM2e{2.0,4.0,6.0}+I{0.2}+G{0.5} | TIM2e+I+G4 | 1 | 2 | 19.230 | 19.195 |
| 69 | DNA | 0/0 | TIM3{2.0,4.0,6.0}+F{0.3,0.1,0.2,0.4} | TIM3+F+ASC | 2 | 6 | 19.211 | 19.229 |
| 70 | DNA | 0/0 | TIM3{2.0,4.0,6.0}+F{0.2,0.1,0.3,0.4}+I{0.2} | TIM3+F+I | 1 | 4 | 21.319 | 20.223 |
| 71 | DNA | 0/0 | TIM3{2.0,4.0,6.0}+F{0.4,0.1,0.2,0.3}+G{0.5} | TIM3+F+G4 | 1 | 4 | 19.340 | 19.439 |
| 72 | DNA | 0/0 | TIM3{2.0,4.0,6.0}+F{0.1,0.2,0.4,0.3}+I{0.2}+G{0.5} | TIM3+F+I+G4 | 1 | 2 | 19.978 | 19.941 |
| 73 | DNA | 0/0 | TIM3e{2.0,4.0,6.0} | TIM3e+ASC | 2 | 4 | 18.704 | 18.744 |
| 74 | DNA | 0/0 | TIM3e{2.0,4.0,6.0}+I{0.2} | TIM3e+I | 1 | 2 | 17.797 | 19.202 |
| 75 | DNA | 0/0 | TIM3e{2.0,4.0,6.0}+G{0.5} | TIM3e+G4 | 1 | 0 | 17.577 | 17.573 |
| 76 | DNA | 0/0 | TIM3e{2.0,4.0,6.0}+I{0.2}+G{0.5} | TIM3e+I+G4 | 1 | 4 | 19.928 | 19.815 |
| 77 | DNA | 0/0 | TVM{2.0,4.0,6.0,8.0}+F{0.3,0.1,0.2,0.4} | TVM+F+ASC | 2 | 0 | 20.090 | 20.103 |
| 78 | DNA | 0/0 | TVM{2.0,4.0,6.0,8.0}+F{0.2,0.1,0.3,0.4}+I{0.2} | TVM+F+I | 1 | 4 | 20.894 | 19.587 |
| 79 | DNA | 0/0 | TVM{2.0,4.0,6.0,8.0}+F{0.4,0.1,0.2,0.3}+G{0.5} | TVM+F+G4 | 1 | 0 | 19.045 | 19.168 |
| 80 | DNA | 0/0 | TVM{2.0,4.0,6.0,8.0}+F{0.1,0.2,0.4,0.3}+I{0.2}+G{0.5} | TVM+F+I+G4 | 1 | 2 | 19.861 | 19.905 |
| 81 | DNA | 0/0 | TVMe{2.0,4.0,6.0,8.0} | TVMe+ASC | 2 | 2 | 21.532 | 21.519 |
| 82 | DNA | 0/0 | TVMe{2.0,4.0,6.0,8.0}+I{0.2} | TVMe+I | 1 | 0 | 20.474 | 20.589 |
| 83 | DNA | 0/0 | TVMe{2.0,4.0,6.0,8.0}+G{0.5} | TVMe+G4 | 1 | 0 | 18.789 | 18.976 |
| 84 | DNA | 0/0 | TVMe{2.0,4.0,6.0,8.0}+I{0.2}+G{0.5} | TVMe+I+G4 | 1 | 6 | 19.383 | 19.419 |
| 85 | DNA | 0/0 | SYM{2.0,4.0,6.0,8.0,10.0} | SYM+ASC | 2 | 0 | 20.516 | 20.523 |
| 86 | DNA | 0/0 | SYM{2.0,4.0,6.0,8.0,10.0}+I{0.2} | SYM+I | 1 | 2 | 20.440 | 20.416 |

| | | | | | | | | |
|---|---|---|---|---|---|---|---|---|
| 87 | DNA | 0/0 | SYM{2.0,4.0,6.0,8.0,10.0}+G{0.5} | SYM+G4 | 1 | 4 | 19.600 | 19.808 |
| 88 | DNA | 0/0 | SYM{2.0,4.0,6.0,8.0,10.0}+I{0.2}+G{0.5} | SYM+I+G4 | 1 | 4 | 19.429 | 19.317 |
| 89 | Protein | 0/0 | Blosum62 | Blosum62 | 1 | 6 | 19.568 | 19.507 |
| 90 | Protein | 0/0 | cpREV | cpREV | 1 | 4 | 16.273 | 16.165 |
| 91 | Protein | 0/0 | Dayhoff | DCMut | 2 | 4 | 22.340 | 22.375 |
| 92 | Protein | 0/0 | DCMut | Dayhoff | 2 | 2 | 21.328 | 21.333 |
| 93 | Protein | 0/0 | FLU | FLU | 1 | 2 | 17.545 | 17.511 |
| 94 | Protein | 0/0 | HIVb | HIVb | 1 | 0 | 20.710 | 20.578 |
| 95 | Protein | 0/0 | HIVw | HIVw | 1 | 6 | 20.923 | 20.782 |
| 96 | Protein | 0/0 | JTT | JTT | 1 | 4 | 18.371 | 18.533 |
| 97 | Protein | 0/0 | JTTDCMut | JTTDCMut | 1 | 12 | 20.420 | 20.741 |
| 98 | Protein | 0/0 | LG | LG | 1 | 2 | 22.356 | 22.245 |
| 99 | Protein | 0/0 | mtART | mtART | 1 | 4 | 17.961 | 18.250 |
| 100 | Protein | 0/0 | mtMAM | mtMAM | 1 | 2 | 18.460 | 18.556 |
| 101 | Protein | 0/0 | mtREV | mtREV | 1 | 8 | 18.682 | 18.737 |
| 102 | Protein | 0/0 | mtZOA | mtZOA | 1 | 12 | 19.678 | 19.724 |
| 103 | Protein | 0/0 | mtMet | mtMet | 1 | 4 | 23.644 | 23.674 |
| 104 | Protein | 0/0 | mtVer | mtVer | 1 | 4 | 17.670 | 17.727 |
| 105 | Protein | 0/0 | mtInv | mtInv | 1 | 6 | 19.786 | 19.849 |
| 106 | Protein | 0/0 | Poisson | Poisson | 1 | 2 | 18.748 | 18.896 |
| 107 | Protein | 0/0 | PMB | PMB | 1 | 4 | 21.342 | 21.231 |
| 108 | Protein | 0/0 | rtREV | rtREV | 1 | 2 | 18.487 | 18.679 |
| 109 | Protein | 0/0 | VT | VT | 1 | 2 | 18.501 | 18.369 |
| 110 | Protein | 0/0 | WAG | WAG | 1 | 2 | 20.125 | 20.276 |

| | | | | | | | | |
|---|---|---|---|---|---|---|---|---|
| 111 | Protein | 0/0 | GTR20+F | GTR20+F | 1 | 2 | 21.327 | 21.343 |
| 112 | Codon | 0/0 | MG{1.0}+F3X4{0.2,0.3,0.4,0.1,0.4,0.3,0.2,0.1,0.1,0.2,0.3,0.4} | MG+F3X4 | 1 | 0 | 18.759 | 18.822 |
| 113 | Codon | 0/0 | MGK{1.0,0.3}+F3X4{0.2,0.3,0.4,0.1,0.4,0.3,0.2,0.1,0.1,0.2,0.3,0.4} | MG+F3X4 | 2 | 0 | 22.478 | 22.445 |
| 114 | Codon | 0/0 | MG1KTS{1.0,0.3}+F3X4{0.2,0.3,0.4,0.1,0.4,0.3,0.2,0.1,0.1,0.2,0.3,0.4} | MG+F3X4 | 3 | 2 | 19.738 | 19.645 |
| 115 | Codon | 0/0 | MG1KTV{1.0,0.3}+F3X4{0.2,0.3,0.4,0.1,0.4,0.3,0.2,0.1,0.1,0.2,0.3,0.4} | MG1KTV+F3X4 | 1 | 6 | 19.373 | 19.238 |
| 116 | Codon | 0/0 | GY{1.0,0.3}+F | GY+F | 1 | 2 | 19.894 | 19.809 |
| 117 | Codon | 0/0 | GY1KTS{1.0,0.3}+F | GY+F | 2 | 0 | 19.881 | 19.746 |
| 118 | Codon | 0/0 | GY1KTV{1.0,0.3}+F | GY1KTV+F | 1 | 0 | 20.767 | 20.937 |
| 119 | Codon | 0/0 | GY2K{1.0,0.3,0.5}+F | GY+F | 2 | 0 | 18.625 | 18.433 |
| 120 | Codon | 0/0 | ECMK07+F3X4{0.2,0.3,0.4,0.1,0.4,0.3,0.2,0.1,0.1,0.2,0.3,0.4}+R3{0.5,2.0,0.2,3.5,0.3,0.5} | KOSI07+F3X4+R4 | 4 | 6 | 25.143 | 19.581 |
| 121 | Codon | 0/0 | ECMrest+F3X4{0.2,0.3,0.4,0.1,0.4,0.3,0.2,0.1,0.1,0.2,0.3,0.4}+R3{0.5,2.0,0.2,3.5,0.3,0.5} | ECMrest+F3X4+R4 | 3 | 6 | 22.503 | 18.949 |
| 122 | Codon | 0/0 | ECMS05+F3X4{0.2,0.3,0.4,0.1,0.4,0.3,0.2,0.1,0.1,0.2,0.3,0.4}+R3{0.5,2.0,0.2,3.5,0.3,0.5} | GY+F3X4+R5 | 8 | 4 | 22.606 | 20.269 |
| 123 | Codon | 0/0 | ECMK07_GY2K{1.0,0.3,0.5}+F+R3{0.5,2.0,0.2,3.5,0.3,0.5} | ECMK07_GY2K+F+R5 | 13 | 2 | 22.528 | 22.401 |
| 124 | Morphological | 0/0 | MK | MK+FQ | 1 | 0 | 19.749 | 19.779 |
| 125 | Morphological | 0/0 | MK+I{0.2} | MK+FQ+I | 1 | 0 | 20.924 | 21.043 |
| 126 | Morphological | 0/0 | MK+G{0.5} | MK+FQ+G4 | 1 | 0 | 18.694 | 18.666 |
| 127 | Morphological | 0/0 | MK+I{0.2}+G{0.5} | MK+FQ+I+G4 | 1 | 0 | 19.503 | 19.407 |
| 128 | Morphological | 0/0 | ORDERED | ORDERED+FQ+ASC | 2 | 0 | 21.046 | 21.035 |

| | | | | | | | | |
|---|---|---|---|---|---|---|---|---|
| 129 | Morphological | 0/0 | ORDERED+I{0.2} | ORDERED+FQ+I | 1 | 2 | 19.734 | 19.803 |
| 130 | Morphological | 0/0 | ORDERED+G{0.5} | ORDERED+FQ+G4 | 1 | 0 | 20.130 | 20.120 |
| 131 | Morphological | 0/0 | ORDERED+I{0.2}+G{0.5} | ORDERED+FQ+I+G4 | 1 | 2 | 18.243 | 18.053 |
| 132 | Binary | 0/0 | JC2 | JC2+FQ+ASC | 2 | 8 | 22.133 | 22.067 |
| 133 | Binary | 0/0 | JC2+I{0.2} | JC2+FQ+I | 1 | 2 | 19.783 | 20.044 |
| 134 | Binary | 0/0 | JC2+G{0.5} | JC2+FQ+G4 | 1 | 2 | 21.888 | 21.809 |
| 135 | Binary | 0/0 | JC2+I{0.2}+G{0.5} | JC2+FQ+I+G4 | 1 | 2 | 20.121 | 20.073 |
| 136 | Binary | 0/0 | GTR2+F{0.7/0.3} | GTR2+FO+ASC | 2 | 4 | 22.119 | 22.103 |
| 137 | Binary | 0/0 | GTR2+F{0.7/0.3}+I{0.2} | GTR2+FO+I | 1 | 12 | 20.829 | 20.797 |
| 138 | Binary | 0/0 | GTR2+F{0.7/0.3}+G{0.5} | GTR2+FO+G4 | 1 | 2 | 20.547 | 20.574 |
| 139 | Binary | 0/0 | GTR2+F{0.7/0.3}+I{0.2}+G{0.5} | GTR2+FO+I+G4 | 1 | 0 | 18.835 | 19.043 |
| Lie Markov models | | | | | | | | |
| 140 | Protein | 0/0 | 1.1 | JC+ASC | 8 | 2 | 19.677 | 19.797 |
| 141 | Protein | 0/0 | RY10.12{0.5,0.6,0.9,0.2,0.1,0.4,0.7,0.8,0.3}+F{0.2/0.3/0.1/0.4} | RY10.12+ASC | 2 | 0 | 18.151 | 18.826 |
| 142 | Protein | 0/0 | WS10.12{-0.5,0.6,0.9,0.2,-0.1,0.4,0.7,-0.8,-0.3}+F{0.2/0.3/0.1/0.4} | UNREST+FO+ASC | 6 | 2 | 18.773 | 19.003 |
| 143 | Protein | 0/0 | MK10.12{0.5,0.6,0.9,0.2,0.1,0.4,0.7,0.8,0.3}+F{0.2/0.3/0.1/0.4} | UNREST+FO+ASC | 4 | 2 | 17.920 | 18.524 |
| 144 | Protein | 0/0 | RY10.34{0.5,-0.6,0.8,0.2,-0.1,0.4,0.7,-0.8,-0.35}+F{0.2/0.3/0.1/0.4} | UNREST+FO+ASC | 4 | 2 | 17.901 | 18.240 |
| 145 | Protein | 0/0 | WS10.34{-0.3,0.15,-0.9,0.2,0.6,-0.4,-0.7,0.8,0.5}+F{0.2/0.3/0.1/0.4} | UNREST+FO+ASC | 6 | 0 | 17.694 | 21.320 |
| 146 | Protein | 0/0 | MK10.34{0.8,-0.5,0.9,-0.35,-0.1,0.4,-0.75,0.6,-0.2}+F{0.2/0.3/0.1/0.4} | UNREST+FO+ASC | 6 | 0 | 21.224 | 21.189 |

| | | | | | | | | |
|---|---|---|---|---|---|---|---|---|
| 147 | Protein | 0/0 | 12.12{0.5,0.6,0.9,0.2,0.1,0.4,0.7,0.8,0.3,0.15,0.65 }+F{0.2/0.3/0.1/0.4} | UNREST+FO+ ASC | 4 | 0 | 19.762 | 19.232 |
| 148 | Protein | 0/0 | RY2.2b{0.2} | K2P+ASC | 12 | 0 | 20.637 | 20.689 |
| 149 | Protein | 0/0 | WS2.2b{0.6} | K3P+ASC | 4 | 2 | 17.860 | 17.887 |
| 150 | Protein | 0/0 | MK2.2b{0.6} | K3P+ASC | 4 | 0 | 20.350 | 20.433 |
| 151 | Protein | 0/0 | 3.3a{0.6,-0.5} | K3P+ASC | 6 | 0 | 19.513 | 19.647 |
| 152 | Protein | 0/0 | RY3.3b{0.6,0.9} | RY3.3b+ASC | 2 | 0 | 19.245 | 19.307 |
| 153 | Protein | 0/0 | WS3.3b{0.6,0.9} | WS3.3b+ASC | 2 | 0 | 20.406 | 20.401 |
| 154 | Protein | 0/0 | MK3.3b{0.6,0.9} | MK3.3b+ASC | 2 | 2 | 21.372 | 21.307 |
| 155 | Protein | 0/0 | RY3.3c{-0.2,0.9} | TNe+ASC | 10 | 2 | 18.676 | 18.710 |
| 156 | Protein | 0/0 | WS3.3c{0.6,0.1} | WS3.3c+ASC | 2 | 0 | 19.210 | 19.181 |
| 157 | Protein | 0/0 | MK3.3c{0.6,0.9} | MK3.3c | 1 | 0 | 21.364 | 21.378 |
| 158 | Protein | 0/0 | RY3.4{-0.6,0.1}+F{0.3} | RY3.4+ASC | 2 | 6 | 20.000 | 19.899 |
| 159 | Protein | 0/0 | WS3.4{-0.6,0.9}+F{0.3} | WS3.4+ASC | 2 | 2 | 19.134 | 19.446 |
| 160 | Protein | 0/0 | MK3.4{0.6,-0.5}+F{0.3} | MK3.4+ASC | 2 | 2 | 20.333 | 21.252 |
| 161 | Protein | 0/0 | 4.4a{0.6,-0.5,0.1}+F{0.2/0.3/0.1/0.4} | UNREST+FO+ ASC | 4 | 2 | 15.938 | 18.585 |
| 162 | Protein | 0/0 | RY4.4b{0.1,-0.5,0.7}+F{0.3} | RY4.4b+ASC | 2 | 2 | 13.972 | 16.428 |
| 163 | Protein | 0/0 | WS4.4b{0.1,-0.5,-0.1}+F{0.3} | WS4.4b+ASC | 2 | 2 | 20.003 | 18.856 |
| 164 | Protein | 0/0 | MK4.4b{-0.6,0.7,0.1}+F{0.3} | MK4.4b+ASC | 2 | 2 | 20.403 | 20.674 |
| 165 | Protein | 0/0 | RY4.5a{0.6,0.9,0.2}+F{0.3} | RY4.5a+ASC | 2 | 0 | 19.763 | 19.721 |
| 166 | Protein | 0/0 | WS4.5a{0.6,0.9,0.2}+F{0.3} | WS4.5a+ASC | 2 | 0 | 21.153 | 21.085 |
| 167 | Protein | 0/0 | MK4.5a{0.6,0.9,0.2}+F{0.3} | MK4.5a+ASC | 2 | 4 | 20.536 | 20.532 |
| 168 | Protein | 0/0 | RY4.5b{0.6,0.9,0.2}+F{0.3} | RY4.5b+ASC | 2 | 2 | 20.246 | 20.290 |
| 169 | Protein | 0/0 | WS4.5b{0.6,0.9,0.2}+F{0.3} | WS4.5b+ASC | 2 | 2 | 21.117 | 21.115 |

| | | | | | | | | |
|---|---|---|---|---|---|---|---|---|
| 170 | Protein | 0/0 | MK4.5b{0.6,-0.9,0.2}+F{0.3} | MK4.5b+ASC | 2 | 0 | 19.579 | 19.562 |
| 171 | Protein | 0/0 | RY5.11a{0.5,-0.6,0.9,-0.25}+F{0.3/0.1} | UNREST+FO+ ASC | 6 | 2 | 20.342 | 20.693 |
| 172 | Protein | 0/0 | WS5.11a{0.5,0.6,0.9,0.2}+F{0.3/0.1} | UNREST+FO+ ASC | 4 | 0 | 19.621 | 20.150 |
| 173 | Protein | 0/0 | MK5.11a{0.5,0.6,0.9,0.2}+F{0.3/0.1} | UNREST+FO+ ASC | 4 | 0 | 19.335 | 19.943 |
| 174 | Protein | 0/0 | RY5.11b{0.5,0.6,0.9,0.2} | RY5.11b+ASC | 2 | 0 | 20.044 | 20.066 |
| 175 | Protein | 0/0 | WS5.11b{0.5,0.6,0.9,0.2} | WS5.11b+ASC | 2 | 0 | 17.502 | 17.481 |
| 176 | Protein | 0/0 | MK5.11b{0.5,0.6,0.9,0.2} | MK5.11b+ASC | 2 | 2 | 23.573 | 23.650 |
| 177 | Protein | 0/0 | RY5.11c{0.5,0.6,0.9,0.2} | RY5.11c+ASC | 2 | 4 | 18.108 | 18.097 |
| 178 | Protein | 0/0 | WS5.11c{0.5,0.6,0.9,0.2} | WS5.11c+ASC | 2 | 0 | 17.608 | 17.614 |
| 179 | Protein | 0/0 | MK5.11c{0.5,0.6,0.9,0.2} | MK5.11c+ASC | 2 | 0 | 19.834 | 19.838 |
| 180 | Protein | 0/0 | RY5.16{0.5,0.6,0.9,0.2}+F{0.3} | RY5.16+ASC | 2 | 0 | 20.847 | 20.799 |
| 181 | Protein | 0/0 | WS5.16{0.5,0.6,0.9,0.2}+F{0.3} | WS5.16+ASC | 2 | 2 | 19.756 | 19.715 |
| 182 | Protein | 0/0 | MK5.16{0.5,0.6,0.9,0.2}+F{0.3} | MK5.16+ASC | 2 | 0 | 19.989 | 20.067 |
| 183 | Protein | 0/0 | RY5.6a{0.5,0.6,0.9,0.2} | RY5.6a+ASC | 2 | 0 | 21.334 | 21.199 |
| 184 | Protein | 0/0 | WS5.6a{0.5,0.6,0.9,0.2} | WS5.6a+ASC | 2 | 0 | 20.376 | 20.483 |
| 185 | Protein | 0/0 | MK5.6a{0.55,-0.6,0.9,0.25} | MK5.6a+ASC | 2 | 0 | 21.753 | 21.814 |
| 186 | Protein | 0/0 | RY5.6b{0.5,-0.6,0.9,-0.2}+F{0.2/0.3/0.1/0.4} | RY5.6b+ASC | 2 | 2 | 20.119 | 20.252 |
| 187 | Protein | 0/0 | WS5.6b{0.15,-0.4,-0.8,0.5}+F{0.2/0.3/0.1/0.4} | UNREST+FO+ ASC | 6 | 2 | 18.836 | 19.335 |
| 188 | Protein | 0/0 | MK5.6b{-0.2,-0.65,0.9,-0.8}+F{0.2/0.3/0.1/0.4} | UNREST+FO+ ASC | 6 | 2 | 18.799 | 19.695 |
| 189 | Protein | 0/0 | RY5.7a{0.5,0.6,0.9,0.2}+F{0.3/0.1} | UNREST+FO+ ASC | 4 | 4 | 17.356 | 17.636 |
| 190 | Protein | 0/0 | WS5.7a{0.5,0.6,0.9,0.2}+F{0.3/0.1} | UNREST+FO+ ASC | 4 | 0 | 18.799 | 19.024 |

| | | | | | | | | |
|---|---|---|---|---|---|---|---|---|
| 191 | Protein | 0/0 | MK5.7a{0.5,0.6,0.9,0.2}+F{0.3/0.1} | UNREST+FO+ ASC | 4 | 6 | 17.740 | 17.957 |
| 192 | Protein | 0/0 | RY5.7b{0.5,0.6,0.9,0.2} | RY5.7b+ASC | 2 | 2 | 21.681 | 21.684 |
| 193 | Protein | 0/0 | WS5.7b{0.5,0.6,0.9,0.2} | WS5.7b+ASC | 2 | 4 | 22.597 | 22.567 |
| 194 | Protein | 0/0 | MK5.7b{0.5,0.6,0.9,0.2} | MK5.7b+ASC | 2 | 0 | 19.275 | 19.289 |
| 195 | Protein | 0/0 | RY5.7c{0.5,0.6,0.9,0.2} | RY5.7c+ASC | 2 | 2 | 23.344 | 23.285 |
| 196 | Protein | 0/0 | WS5.7c{0.5,0.6,0.9,0.2} | WS5.7c+ASC | 2 | 2 | 20.398 | 20.446 |
| 197 | Protein | 0/0 | MK5.7c{0.5,0.6,0.9,0.2} | MK5.7c+ASC | 2 | 2 | 19.571 | 19.605 |
| 198 | Protein | 0/0 | RY6.17a{0.5,0.6,0.9,0.2,0.1}+F{0.3} | RY6.17a+ASC | 2 | 2 | 19.157 | 19.206 |
| 199 | Protein | 0/0 | WS6.17a{0.5,-0.6,-0.7,0.2,-0.1}+F{0.3} | WS6.17a+ASC | 2 | 0 | 21.618 | 21.650 |
| 200 | Protein | 0/0 | MK6.17a{0.5,0.6,0.9,0.2,0.1}+F{0.3} | MK6.17a+ASC | 2 | 2 | 23.593 | 23.640 |
| 201 | Protein | 0/0 | RY6.17b{0.5,0.6,0.9,0.2,0.1}+F{0.3} | RY6.17b+ASC | 2 | 4 | 17.987 | 17.911 |
| 202 | Protein | 0/0 | WS6.17b{0.5,0.6,0.9,0.2,0.1}+F{0.3} | WS6.17b+ASC | 2 | 4 | 23.100 | 23.024 |
| 203 | Protein | 0/0 | MK6.17b{0.5,0.6,0.9,0.2,0.1}+F{0.3} | MK6.17b+ASC | 2 | 0 | 21.355 | 21.281 |
| 204 | Protein | 0/0 | RY6.6{0.5,0.6,0.9,0.2,0.1}+F{0.3} | RY6.6+ASC | 2 | 2 | 18.793 | 18.712 |
| 205 | Protein | 0/0 | WS6.6{0.5,0.6,0.9,0.2,0.1}+F{0.3} | WS6.6+ASC | 2 | 0 | 18.756 | 18.794 |
| 206 | Protein | 0/0 | MK6.6{0.5,0.7,0.9,0.3,0.1}+F{0.3} | MK6.6+ASC | 2 | 4 | 21.716 | 21.806 |
| 207 | Protein | 0/0 | 6.7a{0.5,-0.65,0.9,-0.25,0.1}+F{0.2/0.3/0.1/0.4} | UNREST+FO+ ASC | 4 | 2 | 19.566 | 21.307 |
| 208 | Protein | 0/0 | RY6.7b{-0.5,0.6,-0.9,0.2,-0.1}+F{0.2/0.3/0.1/0.4} | RY6.7b+ASC | 2 | 2 | 16.927 | 16.619 |
| 209 | Protein | 0/0 | WS6.7b{0.5,0.6,0.9,0.2,0.1}+F{0.2/0.3/0.1/0.4} | UNREST+FO+ ASC | 4 | 0 | 19.180 | 19.723 |
| 210 | Protein | 0/0 | MK6.7b{0.55,-0.4,0.9,- 0.2,0.1}+F{0.2/0.3/0.1/0.4} | UNREST+FO+ ASC | 6 | 0 | 16.825 | 17.920 |
| 211 | Protein | 0/0 | RY6.8a{0.5,-0.6,0.7,0.2,-0.15}+F{0.2/0.3/0.1/0.4} | RY6.8a+ASC | 2 | 2 | 15.825 | 18.221 |
| 212 | Protein | 0/0 | WS6.8a{0.5,0.6,0.9,0.2,0.1}+F{0.2/0.3/0.1/0.4} | WS6.8a+ASC | 2 | 8 | 17.771 | 18.188 |

| | | | | | | | | |
|---|---|---|---|---|---|---|---|---|
| 213 | Protein | 0/0 | MK6.8a{0.5,0.6,0.9,0.2,0.1}+F{0.2/0.3/0.1/0.4} | UNREST+FO+ ASC | 4 | 0 | 19.855 | 20.618 |
| 214 | Protein | 0/0 | RY6.8b{0.5,0.6,0.9,0.2,0.1}+F{0.3} | RY6.8b+ASC | 2 | 0 | 17.673 | 17.850 |
| 215 | Protein | 0/0 | WS6.8b{0.5,0.6,0.9,0.2,0.1}+F{0.3} | WS6.8b+ASC | 2 | 0 | 19.355 | 19.574 |
| 216 | Protein | 0/0 | MK6.8b{0.5,0.6,0.9,0.2,0.1}+F{0.3} | MK6.8b+ASC | 2 | 0 | 18.698 | 18.988 |
| 217 | Protein | 0/0 | RY8.10a{0.5,-0.6,-0.9,-0.2,0.1,0.4,0.7}+F{0.2/0.3/0.1/0.4} | RY8.10a+ASC | 2 | 2 | 16.952 | 17.422 |
| 218 | Protein | 0/0 | WS8.10a{0.5,0.6,0.9,0.2,0.1,0.4,0.7}+F{0.2/0.3/0.1/0.4} | UNREST+FO+ ASC | 4 | 2 | 19.493 | 19.560 |
| 219 | Protein | 0/0 | MK8.10a{0.5,0.6,0.9,0.2,0.1,0.4,0.7}+F{0.2/0.3/0.1/0.4} | UNREST+FO+ ASC | 4 | 4 | 20.861 | 21.537 |
| 220 | Protein | 0/0 | RY8.10b{0.5,0.6,0.9,0.2,0.1,0.4,0.7}+F{0.3} | RY8.10b+ASC | 2 | 4 | 20.006 | 20.449 |
| 221 | Protein | 0/0 | WS8.10b{0.5,0.6,0.9,0.2,0.1,0.4,0.7}+F{0.3} | WS8.10b+ASC | 2 | 0 | 18.852 | 19.275 |
| 222 | Protein | 0/0 | MK8.10b{0.5,0.6,0.9,0.2,0.1,0.4,0.7}+F{0.3} | MK8.10b+ASC | 2 | 0 | 19.882 | 20.367 |
| 223 | Protein | 0/0 | RY8.16{0.5,0.6,0.9,0.2,0.1,0.4,0.7}+F{0.2/0.3/0.1/0.4} | UNREST+FO+ ASC | 4 | 4 | 21.246 | 20.620 |
| 224 | Protein | 0/0 | WS8.16{0.5,0.6,0.9,0.2,0.1,0.4,0.7}+F{0.2/0.3/0.1/0.4} | WS8.16+ASC | 2 | 0 | 18.289 | 19.226 |
| 225 | Protein | 0/0 | MK8.16{-0.55,0.6,0.8,-0.2,0.1,0.4,-0.7}+F{0.2/0.3/0.1/0.4} | MK8.16+ASC | 2 | 4 | 20.197 | 20.346 |
| 226 | Protein | 0/0 | RY8.17{0.2,-0.6,0.9,0.5,-0.1,0.4,0.7}+F{0.2/0.3/0.1/0.4} | UNREST+FO+ ASC | 6 | 4 | 17.220 | 17.000 |
| 227 | Protein | 0/0 | WS8.17{0.5,0.6,0.9,0.2,0.1,0.4,0.7}+F{0.2/0.3/0.1/0.4} | WS8.17+ASC | 2 | 2 | 18.646 | 19.195 |
| 228 | Protein | 0/0 | MK8.17{0.3,0.6,-0.9,0.2,-0.4,0.4,0.7}+F{0.2/0.3/0.1/0.4} | UNREST+FO+ ASC | 8 | 2 | 19.098 | 19.620 |
| 229 | Protein | 0/0 | RY8.18{0.4,-0.6,0.9,-0.2,0.1,-0.5,0.7}+F{0.2/0.3/0.1/0.4} | UNREST+FO+ ASC | 4 | 0 | 20.931 | 21.249 |
| 230 | Protein | 0/0 | WS8.18{0.5,0.6,0.9,0.2,0.1,0.4,0.7}+F{0.2/0.3/0.1/0.4} | WS8.18+ASC | 2 | 2 | 21.260 | 22.280 |

| | | | | | | | | |
|---|---|---|---|---|---|---|---|---|
| 231 | Protein | 0/0 | MK8.18{0.5,0.85,-0.9,0.2,-0.15,0.4,-0.7}+F{0.2/0.3/0.1/0.4} | MK8.18+ASC | 2 | 0 | 19.306 | 18.755 |
| 232 | Protein | 0/0 | RY8.8{0.5,0.6,0.9,0.2,0.1,0.4,0.7}+F{0.2/0.3/0.1/0.4} | RY8.8+ASC | 2 | 2 | 21.408 | 20.938 |
| 233 | Protein | 0/0 | WS8.8{-0.5,0.6,-0.9,0.2,-0.1,0.4,-0.7}+F{0.2/0.3/0.1/0.4} | UNREST+FO+ASC | 4 | 0 | 18.500 | 18.814 |
| 234 | Protein | 0/0 | MK8.8{0.45,-0.6,0.8,0.3,-0.1,-0.3,-0.7}+F{0.2/0.3/0.1/0.4} | MK8.8+ASC | 2 | 2 | 21.407 | 20.801 |
| 235 | Protein | 0/0 | RY9.20a{0.9,-0.65,0.9,0.2,-0.1,0.4,0.7,-0.8}+F{0.3/0.1} | RY9.20a+ASC | 2 | 2 | 20.044 | 19.519 |
| 236 | Protein | 0/0 | WS9.20a{0.35,0.6,-0.9,0.2,0.1,-0.4,0.7,0.8}+F{0.3/0.1} | UNREST+FO+ASC | 6 | 0 | 16.002 | 16.896 |
| 237 | Protein | 0/0 | MK9.20a{0.5,0.6,0.9,0.2,0.1,0.4,0.7,0.8}+F{0.3/0.1} | MK9.20a+ASC | 2 | 0 | 19.639 | 19.915 |
| 238 | Protein | 0/0 | 9.20b{0.5,0.6,0.9,0.2,0.1,0.4,0.7,0.8} | 9.20b+ASC | 2 | 4 | 20.319 | 20.329 |
| Mixture models | | | | | | | | |
| 239 | Protein | 0/0 | C10 | C10 | 1 | 0 | 19.164 | 19.055 |
| 240 | Protein | 0/0 | EX2 | GTR20+F+R2 | 2 | 2 | 22.741 | 22.874 |
| 241 | Protein | 0/0 | EX3 | EX3+R2 | 5 | 0 | 22.118 | 21.008 |
| 242 | Protein | 0/0 | EHO | EHO | 1 | 0 | 18.576 | 18.627 |
| 243 | Protein | 0/0 | UL2 | UL2+R2 | 10 | 0 | 25.265 | 20.933 |
| 244 | Protein | 0/0 | UL3 | UL3+R2 | 5 | 0 | 22.850 | 20.000 |
| 245 | Protein | 0/0 | EX_EHO | EX_EHO+R2 | 9 | 0 | 22.013 | 19.726 |
| 246 | Protein | 0/0 | LG4M | LG4M | 1 | 4 | 17.510 | 17.523 |
| 247 | Protein | 0/0 | CF4 | CF4+F+G | 4 | 0 | 20.496 | 20.772 |
| 248 | Protein | 0/0 | JTT+CF4+G4 | JTT+CF4+F+G | 3 | 0 | 20.777 | 20.209 |
| 249 | DNA | 0/0 | MIX{JC,HKY{2.0}}+G4 | MIX{JC,HKY+F}+G4 | 1 | 2 | 17.891 | 17.863 |

| | | | | | | | | |
|---|---|---|---|---|---|---|---|---|
| Heterotachy models | | | | | | | | |
| 250 | DNA | 0/0 | GTR{2/3/4/5/6}+F{0.2/0.3/0.1/0.4}+H4{0.15/0.2/0.35/0.3} | GTR+FU+H4 | 1 | 0 | 21.524 | 21.599 |
| 251 | DNA | 0/0 | MIX{GTR{2/3/4/5/6},GTR{6/2/5/4/7},GTR{4/7/6/2/8},GTR{5/2/7/6/9}}+F{0.2/0.3/0.1/0.4}*H4{0.15/0.2/0.35/0.3} | GTR*H4 | 1 | 0 | 26.843 | 26.872 |
| 252 | DNA | 0/0 | MIX{GTR{2/3/4/5/6}+F{0.2/0.3/0.4/0.1},GTR{6/2/5/4/7}+F{0.3/0.2/0.1/0.4},GTR{4/7/6/2/8}+F{0.4/0.1/0.3/0.2},GTR{5/2/7/6/9}+F{0.1/0.2/0.4/0.3}}*H4{0.15/0.2/0.35/0.3} | GTR+FO*H4 | 1 | 0 | 20.897 | 20.902 |
| Insertion-deletion (Indel) models | | | | | | | | |
| 253 | DNA | 0/0.02 | GTR{1.658853/3.559765/0.056937/1.192477/0.329238}+F{0.386134/0.22998/0.174036/0.20985} | GTR+F | 1 | 2 | 21.347 | 21.385 |
| 254 | DNA | 0/0.04 | GTR{1.32776/8.778905/0.84919/0.515804/1.901472}+F{0.172235/0.264059/0.264182/0.299524} | GTR+F | 1 | 0 | 19.780 | 19.890 |
| 255 | DNA | 0/0.06 | GTR{1.119447/3.559765/0.221697/1.014989/20.711653}+F{0.305724/0.243384/0.196665/0.254226} | GTR+F+G4 | 2 | 0 | 20.064 | 19.769 |
| 256 | DNA | 0/0.08 | GTR{0.086494/2.323423/2.809304/1.318319/4.171516}+F{0.284127/0.219497/0.186409/0.309967} | GTR+F | 1 | 2 | 20.083 | 20.194 |
| 257 | DNA | 0/0.1 | GTR{2.685037/24.868964/1.567923/0.101501/3.578998}+F{0.266817/0.202613/0.308702/0.221868} | GTR+F | 1 | 0 | 19.517 | 19.706 |
| 258 | DNA | 0.02/0 | GTR{8.193704/8.778905/3.613696/0.163775/31.523798}+F{0.292788/0.20068/0.175349/0.331184} | GTR+F | 1 | 0 | 21.322 | 21.315 |
| 259 | DNA | 0.02/0.02 | GTR{0.202229/4.955478/1.841897/1.864075/0.222629}+F{0.303056/0.194328/0.159597/0.343019} | GTR+F | 1 | 2 | 21.568 | 21.543 |
| 260 | DNA | 0.02/0.04 | GTR{8.193704/2.035624/0.340067/1.636181/214.150146}+F{0.273051/0.240699/0.189678/0.296573} | GTR+F | 1 | 0 | 18.628 | 18.706 |
| 261 | DNA | 0.02/0.06 | GTR{1.225831/0.318379/1.382544/5.206825/4.826486}+F{0.329796/0.339286/0.131282/0.199636} | GTR+F | 1 | 2 | 20.093 | 20.150 |

| | | | | | | | | |
|---|---|---|---|---|---|---|---|---|
| 262 | DNA | 0.02/0.08 | GTR{2.666457/4.112417/0.177203/0.910549/10.179543}+F{0.243052/0.364955/0.187378/0.204614} | GTR+F | 1 | 0 | 19.308 | 19.341 |
| 263 | DNA | 0.02/0.1 | GTR{8.193704/7.74051/1.841897/0.841534/1.788653}+F{0.28858/0.320342/0.182192/0.208886} | GTR+F | 1 | 2 | 19.756 | 19.811 |
| 264 | DNA | 0.04/0 | GTR{1.259178/14.491652/0.84919/1.192477/17.608509}+F{0.364391/0.166043/0.209958/0.259609} | GTR+F | 1 | 0 | 18.784 | 18.877 |
| 265 | DNA | 0.04/0.02 | GTR{1.622091/3.548607/2.64403/0.902628/3.029666}+F{0.293661/0.223556/0.163246/0.319537} | GTR+F | 1 | 0 | 21.226 | 21.255 |
| 266 | DNA | 0.04/0.04 | GTR{1.841625/5.32596/2.941686/0.515804/5.660189}+F{0.134343/0.270785/0.293569/0.301302} | GTR+F | 1 | 4 | 17.389 | 17.401 |
| 267 | DNA | 0.04/0.06 | GTR{1.119447/7.159051/2.006633/0.975949/0.970356}+F{0.341057/0.27335/0.177055/0.208539} | GTR+F | 1 | 4 | 19.933 | 19.986 |
| 268 | DNA | 0.04/0.08 | GTR{2.379396/4.955478/0.989033/13.917175/10.179543}+F{0.216181/0.363212/0.16473/0.255877} | GTR+F | 1 | 4 | 20.559 | 20.546 |
| 269 | DNA | 0.04/0.1 | GTR{1.487735/1.53548/1.741463/1.695331/11.237555}+F{0.335028/0.192155/0.194448/0.278368} | GTR+F | 1 | 2 | 22.209 | 22.037 |
| 270 | DNA | 0.06/0 | GTR{1.492042/1.529111/0.876352/0.781435/9.382395}+F{0.320038/0.20406/0.201034/0.274867} | GTR+F | 1 | 4 | 21.561 | 21.586 |
| 271 | DNA | 0.06/0.02 | GTR{2.666457/4.955478/0.221697/1.521687/8.655391}+F{0.314747/0.227549/0.20465/0.253054} | GTR+F | 1 | 0 | 18.015 | 18.032 |
| 272 | DNA | 0.06/0.04 | GTR{1.119447/2.949316/0.478944/0.517493/17.483976}+F{0.357602/0.191959/0.165626/0.284813} | GTR+F | 1 | 2 | 20.911 | 21.021 |
| 273 | DNA | 0.06/0.06 | GTR{2.857435/9.196923/0.351809/4.999912/1.653673}+F{0.303202/0.195056/0.184061/0.317681} | GTR+F | 1 | 0 | 20.761 | 20.676 |
| 274 | DNA | 0.06/0.08 | GTR{2.857435/7.236792/0.525951/0.783632/5.026411}+F{0.23663/0.161377/0.376/0.225992} | GTR+F | 1 | 4 | 20.621 | 20.645 |
| 275 | DNA | 0.06/0.1 | GTR{0.346774/4.384402/2.905711/3.191284/8.655391}+F{0.329501/0.271937/0.21811/0.180452} | GTR+F | 1 | 2 | 20.076 | 20.025 |

| | | | | | | | | |
|---|---|---|---|---|---|---|---|---|
| 276 | DNA | 0.08/0 | GTR{0.918148/7.28152/0.112665/0.975949/8.452647}+F{0.263484/0.318991/0.229091/0.188434} | GTR+F | 1 | 2 | 21.971 | 21.939 |
| 277 | DNA | 0.08/0.02 | GTR{3.806577/8.494684/1.094147/3.506839/3.227558}+F{0.38109/0.174324/0.22796/0.216625} | GTR+F | 1 | 0 | 20.419 | 20.511 |
| 278 | DNA | 0.08/0.04 | GTR{1.436101/5.051975/0.112665/0.930071/7.797079}+F{0.269989/0.245349/0.261474/0.223188} | GTR+F | 1 | 0 | 18.967 | 19.082 |
| 279 | DNA | 0.08/0.06 | GTR{0.389424/2.591557/0.546466/1.288688/20.711653}+F{0.340814/0.169713/0.160952/0.328521} | GTR+F | 1 | 4 | 18.937 | 18.843 |
| 280 | DNA | 0.08/0.08 | GTR{3.166275/24.868964/0.221697/0.470818/3.128172}+F{0.179181/0.20477/0.238503/0.377545} | GTR+F | 1 | 4 | 22.092 | 21.728 |
| 281 | DNA | 0.08/0.1 | GTR{3.049122/8.778905/0.654595/0.586987/0.329238}+F{0.380216/0.183774/0.164769/0.271242} | GTR+F | 1 | 0 | 20.349 | 20.325 |
| 282 | DNA | 0.1/0 | GTR{1.841625/7.236792/0.417861/1.099721/3.27437}+F{0.311121/0.268752/0.256148/0.163979} | GTR+F | 1 | 2 | 20.760 | 20.755 |
| 283 | DNA | 0.1/0.02 | GTR{5.08016/8.803904/2.809304/2.939371/2.492354}+F{0.298449/0.155919/0.216434/0.329199} | GTR+F | 1 | 2 | 18.955 | 18.966 |
| 284 | DNA | 0.1/0.04 | GTR{2.857435/3.344601/0.876352/2.455665/70.026968}+F{0.253814/0.204526/0.240511/0.301149} | GTR+F | 1 | 0 | 21.113 | 21.135 |
| 285 | DNA | 0.1/0.06 | GTR{5.08016/1.692792/0.431923/2.769857/8.655391}+F{0.379775/0.204224/0.160545/0.255456} | GTR+F | 1 | 0 | 18.434 | 18.450 |
| 286 | DNA | 0.1/0.08 | GTR{1.32776/7.185683/0.567838/13.917175/94.907071}+F{0.273149/0.350457/0.137704/0.238689} | GTR+F | 1 | 2 | 20.981 | 21.044 |
| 287 | DNA | 0.1/0.1 | GTR{0.975527/7.153133/0.59506/11.3205/10.321999}+F{0.240426/0.200113/0.322616/0.236845} | GTR+F | 1 | 2 | 19.049 | 19.023 |

NOTE.-We simulated 287 MSAs from random 100-tip trees with sequence length of 10K sites (for DNA, binary, morphological, codon data) and 1K sites (for protein data) across a wide range of substitution models and insertion-deletion rates of 0.0, 0.02, 0.04, 0.06, 0.08 and 0.1. These choices of indel rates follow empirical studies (Cartwright 2009). We then ran IQ-TREE to determine the best-fit model using ModelFinder

(Kalyaanamoorthy et al. 2017) and reconstructed phylogenetic trees under the best-fit model. We compared the topology between the true trees and the inferred trees using the Robinson-Foulds distance (Robinson and Foulds 1981). The table shows detailed results, each row corresponds to one test case. In 147 tests (51.22%), the true model was recovered as the best-fit model. In 243 tests (84.67%), 246 tests (85.71%), and 267 tests (93.03%), the true models appear in the top-2, top-3, and top-4 best models, respectively. The average Robinson-Foulds distance between the true trees and the inferred trees across all test cases was 1.99 (s.e. 0.133). That means the inferred trees differed from the true trees by only 0.995 out of 97 (1.03%) internal branches. The tree lengths (sum of branch lengths) of the inferred trees differed from the true trees by only 1.9%.

**Table S2.** Comparison results between AliSim and INDELible on simulating 35 MSAs with different combinations of insertion and deletion rates of 0.0, 0.02, 0.04, 0.06, 0.08 and 0.1.

| Indel-rate | The true (input) model | Alignment length | | | Proportion of gaps (%) | | |
|---|---|---|---|---|---|---|---|
| | | AliSim | INDELible | Difference | AliSim | INDELible | Difference |
| 0/0.02 | GTR{1.658853/3.559765/0.056937/1.192477/0.329238}+F{0.386134/0.22998/0.174036/0.20985} | 10000 | 10000 | 0.00% | 2.73% | 2.74% | 0.01% |
| 0/0.04 | GTR{1.32776/8.778905/0.84919/0.515804/1.901472}+F{0.172235/0.264059/0.264182/0.299524} | 10000 | 10000 | 0.00% | 5.50% | 5.88% | 0.38% |
| 0/0.06 | GTR{1.119447/3.559765/0.221697/1.014989/20.711653}+F{0.305724/0.243384/0.196665/0.254226} | 10000 | 10000 | 0.00% | 7.17% | 7.36% | 0.19% |
| 0/0.08 | GTR{0.086494/2.323423/2.809304/1.318319/4.171516}+F{0.284127/0.219497/0.186409/0.309967} | 10000 | 10000 | 0.00% | 13.47% | 13.55% | 0.08% |
| 0/0.1 | GTR{2.685037/24.868964/1.567923/0.101501/3.578998}+F{0.266817/0.202613/0.308702/0.221868} | 10000 | 10000 | 0.00% | 10.55% | 10.84% | 0.29% |
| 0.02/0 | GTR{8.193704/8.778905/3.613696/0.163775/31.523798}+F{0.292788/0.20068/0.175349/0.331184} | 15583 | 15593 | 0.06% | 32.94% | 32.95% | 0.01% |
| 0.02/0.02 | GTR{0.202229/4.955478/1.841897/1.864075/0.222629}+F{0.303056/0.194328/0.159597/0.343019} | 15311 | 15272 | 0.26% | 34.77% | 34.53% | 0.24% |

| | | | | | | | |
|---|---|---|---|---|---|---|---|
| 0.02/0.04 | GTR{8.193704/2.035624/0.340067/1.636181/214.150146}+F{0.273051/0.240699/0.189678/0.296573} | 14450 | 14459 | 0.06% | 33.64% | 33.36% | 0.28% |
| 0.02/0.06 | GTR{1.225831/0.318379/1.382544/5.206825/4.826486}+F{0.329796/0.339286/0.131282/0.199636} | 14699 | 14688 | 0.07% | 36.81% | 36.40% | 0.41% |
| 0.02/0.08 | GTR{2.666457/4.112417/0.177203/0.910549/10.179543}+F{0.243052/0.364955/0.187378/0.204614} | 14594 | 14416 | 1.23% | 36.09% | 34.89% | 1.20% |
| 0.02/0.1 | GTR{8.193704/7.74051/1.841897/0.841534/1.788653}+F{0.28858/0.320342/0.182192/0.208886} | 14344 | 14384 | 0.28% | 38.12% | 37.94% | 0.18% |
| 0.04/0 | GTR{1.259178/14.491652/0.84919/1.192477/17.608509}+F{0.364391/0.166043/0.209958/0.259609} | 19698 | 19777 | 0.40% | 47.66% | 47.88% | 0.22% |
| 0.04/0.02 | GTR{1.622091/3.548607/2.64403/0.902628/3.029666}+F{0.293661/0.223556/0.163246/0.319537} | 21082 | 20891 | 0.91% | 50.75% | 50.44% | 0.31% |
| 0.04/0.04 | GTR{1.841625/5.32596/2.941686/0.515804/5.660189}+F{0.134343/0.270785/0.293569/0.301302} | 18723 | 18607 | 0.62% | 46.91% | 46.28% | 0.63% |
| 0.04/0.06 | GTR{1.119447/7.159051/2.006633/0.975949/0.970356}+F{0.341057/0.27335/0.177055/0.208539} | 19686 | 19830 | 0.73% | 50.18% | 50.64% | 0.46% |
| 0.04/0.08 | GTR{2.379396/4.955478/0.989033/13.917175/10.179543}+F{0.216181/0.363212/0.16473/0.255877} | 19490 | 19793 | 1.53% | 51.77% | 51.95% | 0.18% |
| 0.04/0.1 | GTR{1.487735/1.53548/1.741463/1.695331/11.237555}+F{0.335028/0.192155/0.194448/0.278368} | 20134 | 20154 | 0.10% | 54.57% | 54.35% | 0.22% |
| 0.06/0 | GTR{1.492042/1.529111/0.876352/0.781435/9.382395}+F{0.320038/0.20406/0.201034/0.274867} | 27483 | 27412 | 0.26% | 60.42% | 60.46% | 0.04% |
| 0.06/0.02 | GTR{2.666457/4.955478/0.221697/1.521687/8.655391}+F{0.314747/0.227549/0.20465/0.253054} | 24769 | 24790 | 0.08% | 56.32% | 56.38% | 0.06% |
| 0.06/0.04 | GTR{1.119447/2.949316/0.478944/0.517493/17.483976}+F{0.357602/0.191959/0.165626/0.284813} | 26136 | 26085 | 0.20% | 60.56% | 60.60% | 0.04% |
| 0.06/0.06 | GTR{2.857435/9.196923/0.351809/4.999912/1.653673}+F{0.303202/0.195056/0.184061/0.317681} | 25211 | 25545 | 1.31% | 60.59% | 60.83% | 0.24% |
| 0.06/0.08 | GTR{2.857435/7.236792/0.525951/0.783632/5.026411}+F{0.23663/0.161377/0.376/0.225992} | 25101 | 25288 | 0.74% | 61.43% | 61.50% | 0.07% |
| 0.06/0.1 | GTR{0.346774/4.384402/2.905711/3.191284/8.655391}+F{0.329501/0.271937/0.21811/0.180452} | 24300 | 24320 | 0.08% | 60.80% | 60.79% | 0.01% |

| | | | | | | | |
|---|---|---|---|---|---|---|---|
| 0.08/0 | GTR{0.918148/7.28152/0.112665/0.975949/8.452647}+F{0.263484/0.318991/0.229091/0.188434} | 34176 | 34407 | 0.67% | 67.02% | 67.41% | 0.39% |
| 0.08/0.02 | GTR{3.806577/8.494684/1.094147/3.506839/3.227558}+F{0.38109/0.174324/0.22796/0.216625} | 32195 | 31698 | 1.57% | 66.30% | 65.85% | 0.45% |
| 0.08/0.04 | GTR{1.436101/5.051975/0.112665/0.930071/7.797079}+F{0.269989/0.245349/0.261474/0.223188} | 29794 | 29675 | 0.40% | 65.12% | 65.11% | 0.01% |
| 0.08/0.06 | GTR{0.389424/2.591557/0.546466/1.288688/20.711653}+F{0.340814/0.169713/0.160952/0.328521} | 29194 | 29016 | 0.61% | 64.99% | 64.65% | 0.34% |
| 0.08/0.08 | GTR{3.166275/24.868964/0.221697/0.470818/3.128172}+F{0.179181/0.20477/0.238503/0.377545} | 31731 | 31654 | 0.24% | 68.43% | 68.46% | 0.03% |
| 0.08/0.1 | GTR{3.049122/8.778905/0.654595/0.586987/0.329238}+F{0.380216/0.183774/0.164769/0.271242} | 29543 | 29899 | 1.19% | 67.07% | 67.26% | 0.19% |
| 0.1/0 | GTR{1.841625/7.236792/0.417861/1.099721/3.27437}+F{0.311121/0.268752/0.256148/0.163979} | 38115 | 38627 | 1.33% | 70.96% | 71.16% | 0.20% |
| 0.1/0.02 | GTR{5.08016/8.803904/2.809304/2.939371/2.492354}+F{0.298449/0.155919/0.216434/0.329199} | 36989 | 36910 | 0.21% | 68.81% | 68.76% | 0.05% |
| 0.1/0.04 | GTR{2.857435/3.344601/0.876352/2.455665/70.026968}+F{0.253814/0.204526/0.240511/0.301149} | 39289 | 39100 | 0.48% | 71.52% | 71.48% | 0.04% |
| 0.1/0.06 | GTR{5.08016/1.692792/0.431923/2.769857/8.655391}+F{0.379775/0.204224/0.160545/0.255456} | 34382 | 34184 | 0.58% | 69.33% | 69.12% | 0.21% |
| 0.1/0.08 | GTR{1.32776/7.185683/0.567838/13.917175/94.907071}+F{0.273149/0.350457/0.137704/0.238689} | 37127 | 36680 | 1.22% | 72.25% | 72.24% | 0.01% |
| 0.1/0.1 | GTR{0.975527/7.153133/0.59506/11.3205/10.321999}+F{0.240426/0.200113/0.322616/0.236845} | 33784 | 33751 | 0.10% | 70.38% | 70.53% | 0.15% |

NOTE.- We simulated 10K-site MSAs from 100-tip trees under the general time-reversible (GTR) model (Tavaré 1986) and used the negative binomial distribution (with a mean of 1.25) to generate the insertion and deletion sizes. Each row shows the alignment length and the proportion of gaps of the simulated MSA for each Indel-rate combination. The average differences in the alignment length and proportion of gaps between MSAs simulated by AliSim and those by INDELible were 0.5% and 0.22%, respectively.

**Table S3.** Switching parameters for different sequence lengths, ranging from 1K to 100K sites with/without continuous rate heterogeneity.

| Model | #Site | Runtime of the Probability matrix approach (ms) | Range of branch length | Switching params |
|---|---|---|---|---|
| GTR+F | 1000 | 342.841 | 0.0001-0.1 | 0.002215 |
| GTR+F | 2000 | 605.015 | 0.0001-0.1 | 0.001097 |
| GTR+F | 3000 | 872.343 | 0.0001-0.1 | 0.000754 |
| GTR+F | 4000 | 1159.497 | 0.0001-0.1 | 0.000579 |
| GTR+F | 5000 | 1425.972 | 0.0001-0.1 | 0.000451 |
| GTR+F | 6000 | 1687.084 | 0.0001-0.1 | 0.000384 |
| GTR+F | 7000 | 1985.005 | 0.0001-0.1 | 0.000335 |
| GTR+F | 8000 | 2271.611 | 0.0001-0.1 | 0.000294 |
| GTR+F | 9000 | 2551.453 | 0.0001-0.1 | 0.000262 |
| GTR+F | 10000 | 2811.463 | 1e-05-0.01 | 0.000232 |
| GTR+F | 20000 | 5666.547 | 1e-05-0.01 | 0.000116 |
| GTR+F | 30000 | 8574.731 | 1e-05-0.01 | 0.000074 |
| GTR+F | 40000 | 11338.240 | 1e-05-0.01 | 0.000052 |
| GTR+F | 50000 | 14101.600 | 1e-05-0.01 | 0.000035 |
| GTR+F | 60000 | 17025.880 | 1e-05-0.01 | 0.000027 |
| GTR+F | 70000 | 19920.400 | 1e-05-0.01 | 0.000026 |
| GTR+F | 80000 | 22567.150 | 1e-05-0.01 | 0.000021 |
| GTR+F | 90000 | 25515.960 | 1e-05-0.01 | 0.000016 |
| GTR+F | 100000 | 28254.480 | 1e-06-0.001 | 0.000014 |
| GTR+F+GC | 1000 | 1433.839 | 0.0001-0.1 | 0.01327 |
| GTR+F+GC | 2000 | 2788.569 | 0.0001-0.1 | 0.006695 |
| GTR+F+GC | 3000 | 4153.132 | 0.0001-0.1 | 0.004471 |
| GTR+F+GC | 4000 | 5506.709 | 0.0001-0.1 | 0.00334 |
| GTR+F+GC | 5000 | 6859.667 | 0.0001-0.1 | 0.002671 |
| GTR+F+GC | 6000 | 8213.643 | 0.0001-0.1 | 0.002245 |
| GTR+F+GC | 7000 | 9573.880 | 0.0001-0.1 | 0.001908 |
| GTR+F+GC | 8000 | 10943.020 | 0.0001-0.1 | 0.001675 |
| GTR+F+GC | 9000 | 12308.660 | 0.0001-0.1 | 0.001472 |

| | | | | |
|---|---|---|---|---|
| GTR+F+GC | 10000 | 13669.790 | 1e-05-0.01 | 0.001328 |
| GTR+F+GC | 20000 | 27292.360 | 1e-05-0.01 | 0.000653 |
| GTR+F+GC | 30000 | 40929.470 | 1e-05-0.01 | 0.000433 |
| GTR+F+GC | 40000 | 54561.300 | 1e-05-0.01 | 0.000301 |
| GTR+F+GC | 50000 | 68394.600 | 1e-05-0.01 | 0.0002 |
| GTR+F+GC | 60000 | 81908.780 | 1e-05-0.01 | 0.000164 |
| GTR+F+GC | 70000 | 95600.800 | 1e-05-0.01 | 0.000144 |
| GTR+F+GC | 80000 | 109123.800 | 1e-05-0.01 | 0.00012 |
| GTR+F+GC | 90000 | 122608.600 | 1e-05-0.01 | 0.000105 |
| GTR+F+GC | 100000 | 136422.700 | 1e-05-0.01 | 0.000091 |

NOTE.- Each row shows the model, the sequence length, the runtime of the probability matrix approach, the range of branch length to apply the binary search, and the switching parameter.

**References**

Cartwright RA. 2009. Problems and solutions for estimating indel rates and length distributions. *Mol Biol Evol*. 26(2):473–480.

De Maio Nicola, Weilguny L, Walker CR, Turakhia Y, Corbett-Detig R, Goldman N. 2021. phastSim: efficient simulation of sequence evolution for pandemic-scale datasets. *bioRxiv*. doi: 10.1101/2021.03.15.435416.

Kalyaanamoorthy S, Minh BQ, Wong TKF, von Haeseler A, Jermiin LS. 2017. ModelFinder: Fast model selection for accurate phylogenetic estimates. *Nat Methods*. 14(6):587–589.

Robinson DF, Foulds LR. 1981. Comparison of phylogenetic trees. *Math Biosci*. 53(1–2):131–147.

Tavaré S. Miura RM 1986. Some probabilistic and statistical problems in the analysis of DNA sequences. *Lect Math life Sci*. 17:57–86.
